# Dorsal peduncular cortex activity modulates affective behavior and fear extinction in mice

**DOI:** 10.1101/2023.04.25.538301

**Authors:** Justin J. Botterill, Abdessattar Khlaifia, Ryan Appings, Jennifer Wilkin, Francesca Violi, Hanista Premachandran, Arely Cruz-Sanchez, Anna Elisabete Canella, Ashutosh Patel, S. Danyal Zaidi, Maithe Arruda-Carvalho

## Abstract

The medial prefrontal cortex (mPFC) is critical to cognitive and emotional function and underlies many neuropsychiatric disorders, including mood, fear and anxiety disorders. In rodents, disruption of mPFC activity affects anxiety- and depression-like behavior, with specialized contributions from its subdivisions. The rodent mPFC is divided into the dorsomedial prefrontal cortex (dmPFC), spanning the anterior cingulate cortex (ACC) and dorsal prelimbic cortex (PL), and the ventromedial prefrontal cortex (vmPFC), which includes the ventral PL, infralimbic cortex (IL), and in some studies the dorsal peduncular cortex (DP) and dorsal tenia tecta (DTT). The DP/DTT have recently been implicated in the regulation of stress- induced sympathetic responses via projections to the hypothalamus. While many studies implicate the PL and IL in anxiety-, depression-like and fear behavior, the contribution of the DP/DTT to affective and emotional behavior remains unknown. Here, we used chemogenetics and optogenetics to bidirectionally modulate DP/DTT activity and examine its effects on affective behaviors, fear and stress responses in C57BL/6J mice. Acute chemogenetic activation of DP/DTT significantly increased anxiety-like behavior in the open field and elevated plus maze tests, as well as passive coping in the tail suspension test. DP/DTT activation also led to an increase in serum corticosterone levels and facilitated auditory fear extinction learning and retrieval. Activation of DP/DTT projections to the dorsomedial hypothalamus (DMH) acutely decreased freezing at baseline and during extinction learning, but did not alter affective behavior. These findings point to the DP/DTT as a new regulator of affective behavior and fear extinction in mice.

## Introduction

The medial prefrontal cortex (mPFC) is a key brain region involved in high-order functions such as attention [1,2], decision making [3,4], working memory [5,6], social behavior [7–10], mood [11,12] and anxiety [7,13,22–30,14–21]. The mPFC integrates inputs from cortical sensory and motor systems, as well as subcortical brain structures [31–36] toward top-down control of output structures such as the amygdala, thalamus, hypothalamus and bed nucleus of the stria terminalis (BNST) [31,34,36–38]. Despite a lack of unified consensus [39], many studies consider the rodent mPFC to be anatomically subdivided into the anterior cingulate cortex (ACC), prelimbic cortex (PL), and the infralimbic cortex (IL) [40], with some dividing the mPFC into two main regions with differences in their connectivity pattern and functional properties [40–43]: the dorsal mPFC (dmPFC), which includes the ACC and dorsal PL, and the ventral mPFC (vmPFC), which comprises the ventral PL, IL and the dorsal peduncular cortex (DP) and dorsal tenia tecta (DTT). Altered mPFC activity underlies many neuropsychiatric disorders, including mood, fear and anxiety disorders [38,44–52]. In rodents, disruption of mPFC activity through lesions, pharmacology, optogenetics or chemogenetics affects anxiety- [29,53–55] and depression-like [56–59] behavior, with evidence supporting specialized contributions from PL and IL subdivisions [21,23–27], as well as their subprojections [58,60]. Similarly, the PL and IL mPFC subdivisions exert distinct roles in auditory fear processing [61–66]. Although there is a considerable body of work implicating the mPFC as a key regulator of affective behavior and emotional learning, most studies focus on its main subdivisions, ACC, PL and IL, leaving subregions such as the DP/DTT understudied.

The DP/DTT is located at the ventral limit of the vmPFC [67], and plays a crucial role in driving thermoregulatory and cardiovascular sympathetic responses during stress [68–70]. The DP/DTT receives inputs from stress and emotion-related brain regions like the thalamus, amygdala, insular cortex and piriform cortex [68–70], and sends glutamatergic excitatory input to the dorsomedial hypothalamus (DMH), a central hub driving sympathetic responses to various stimuli, including psychosocial stress [68,70]. Interestingly, while dmPFC activation abrogates stress-related autonomic and neuroendocrine responses [71–74], DP/DTT activation mimics sympathetic responses to psychosocial stress [68], suggesting a distinct role for the DP/DTT in driving sympathetic stress responses. Given the high prevalence of sympathetic system impairments in anxiety disorders [75–78] and the well-established role of the mPFC in modulating anxiety-, depression-like and cognitive behavior in rodents, we investigated the contribution of the DP/DTT to affective behavior and emotional learning in mice. Our results show that chemogenetic activation of DP/DTT with Designer Receptors Exclusively Activated by Designer Drugs (DREADDs) in C57BL/6J mice increases anxiety-like behaviors in the open field (OFT) and elevated plus maze (EPM) tests, passive coping in the tail suspension test (TST) and facilitates fear extinction learning and extinction retrieval. DP/DTT activation was accompanied by an increase in serum corticosterone levels. Activation of DP/DTT projections to the dorsomedial hypothalamus (DMH) acutely suppressed freezing. These findings uncover a novel role for the DP/DTT in promoting anxiety-like and fear extinction behaviors in mice.

## Results

To explore the contribution of the DP/DTT to anxiety-, depression-like and fear behavior in mice, we first bidirectionally modulated DP/DTT activity across a series of behavioral tasks with excitatory and inhibitory DREADDs. AAV vectors encoding excitatory (hM3D), inhibitory (hM4D) DREADDs or the control fluorophore mCherry were injected into the DP/DTT of C57BL/6J mice. Viral expression was strongly localized to the DP/DTT across each treatment condition (**Supplementary Figure S1**).

### Chemogenetic activation of DP/DTT increases anxiety-like behavior

We first examined the impact of manipulating DP/DTT activity on anxiety-like behavior in freely behaving mice in the OFT (**Figure 1A-F; Supplementary Figure S2A-B**). Mice expressing hM3D, hM4D or mCherry constructs in the DP/DTT region were injected with C21 one hour prior to OF testing (**Figure 1A-B**). Chemogenetic activation of DP/DTT neurons (hM3D mice) decreased the time spent in the center zone of the OF arena compared to mCherry and hM4D mice (**Figure 1C**; One-way ANOVA, F_2,38_ =7.795, *p*=0.002, Tukey’s post hoc tests, hM3D versus mCherry *p*=0.002; hM3D versus hM4D *p*=0.011; **Supplementary Figure S2B**). In contrast, hM3D mice spent significantly more time in the outer zone of the open field apparatus compared to both mCherry control and hM4D mice (**Figure 1D**, one way ANOVA, *F*_2,38_=7.283, *p*=0.002; Tukey’s post hoc tests, hM3D versus mCherry, *p*=0.007 and hM3D versus hM4D, *p*=0.006). Furthermore, the total distance travelled was similar across groups (**Figure 1E**, one way ANOVA, *F*_2,38_=2.292, *p*=0.115), suggesting that the effect of C21-mediated activation of DP/DTT neurons on anxiety-like behavior was not driven by changes in locomotor activity levels. These data show that chemogenetic activation of the DP/DTT induces anxiety-like behavior in the OFT (**Figure 1F**).

**Figure 1.**
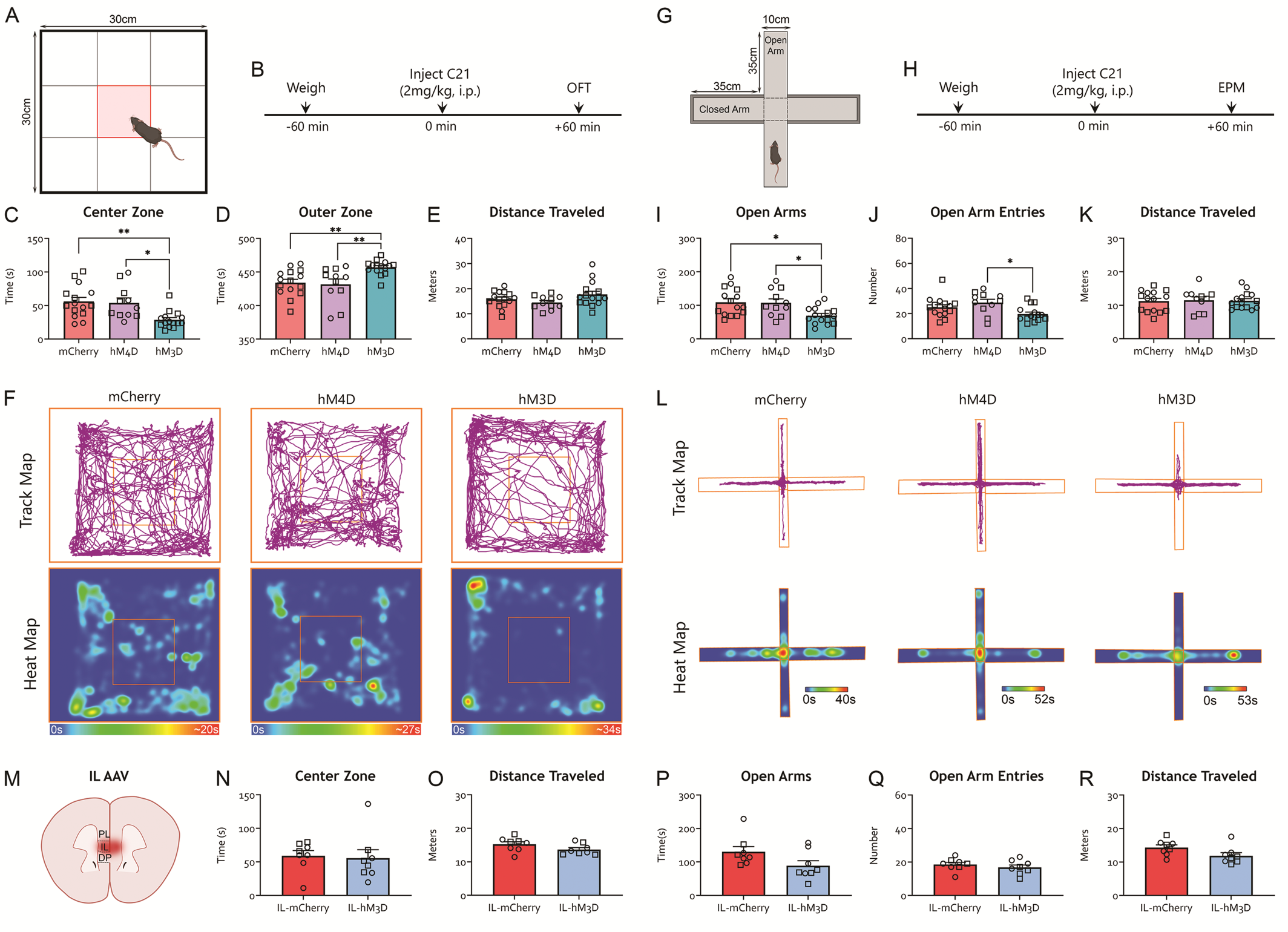
Chemogenetic activation of the DP/DTT increases anxiety-like behavior in the open field and elevated plus maze tests. **A-B.** Experimental timeline for the OFT. Mice previously injected with hM3D, hM4D or control mCherry constructs in the DP/DTT were injected with the DREADD agonist C21 one hour prior to being placed in an open field. The time spent in the center zone (red) and the outer zone (white) was measured. **C.** The time spent in the center zone of the open field was significantly reduced in hM3D mice compared to mCherry and hM4D mice (no difference between mCherry and hM4D, *p*=0.965). **D.** The amount of time spent in the outer zone of the open field arena was significantly greater in the hM3D mice compared to mCherry and hM4D mice (mCherry versus hM4D, *p*=0.938). **E.** The total distance travelled during the test did not differ between groups (one-way ANOVA, *F*_2,38_=2.292, *p*=0.115). **F.** Representative track and heat maps. **G-H.** Experimental timeline for the EPM test. Mice previously injected with hM3D, hM4D or control mCherry constructs in the DP/DTT were injected with the DREADD agonist C21 one hour prior to being placed in an elevated plus maze. The time spent in the open arms vs closed arms was measured. **I.** The time spent in the open arms of the elevated plus maze was significantly reduced in hM3D mice compared to mCherry and hM4D mice (no difference between mCherry and hM4D mice, *p*=0.993). **J.** The number of open arm entries was significantly reduced in hM3D mice compared to hM4D mice (no difference between hM3D and mCherry, *p*=0.136; or hM4D and mCherry, *p*=0.430). **K.** The total distance travelled during the test did not differ between groups (one-way ANOVA, *F*_2,38_=0.022, *p*=0.98). While we found that hM4D males travelled less than hM4D females (see results), separation of the data by sex did not reveal group differences (One-way ANOVA, females F_2,16_= 0.3571, p= 0.7051; males F_2,19_= 0.093, p= 0.9117). **L.** Representative track and heat maps. **M**. Schematic of IL viral targeting. Mice previously injected with hM3D or control mCherry constructs into the IL were injected with the DREADD agonist C21 one hour prior to undergoing the OFT (**N-O**) or EPM test (**P-R**). **N**. IL-hM3D mice spent the same amount of time in the center zone of the OFT as IL-mCherry mice (*t_14_*=0.2502, *p*=0.8061). **O**. OFT distance travelled did not differ between groups (*t_14_*=1.689, *p*=0.1134). **P**. IL-hM3D mice spent the same amount of time in the open arms of the EPM as IL-mCherry mice (*t_14_*=1.916, *p*=0.076). **Q**. IL-hM3D and IL-mCherry mice displayed the same number of open arm entries in the EPM (*t_14_*=0.8721, *p*=0.3979). **R**. EPM distance travelled did not differ between groups (*t_14_*=1.999, *p*=0.0654). *p<0.05, **p<0.01. Male (square) and female (circle) individual datapoints are displayed for transparency (see text for details).

Next, we examined the effects of manipulating DP/DTT activity on anxiety-like behavior using the EPM test (**Figure 1G-L; Supplementary Figure S2C-F**). Consistent with the anxiogenic effect of DP/DTT activation observed in the OFT, hM3D mice spent less time in the open arms of the EPM apparatus compared to mCherry control and hM4D mice (**Figure 1I**; one-way ANOVA, *F*_2,38_=5.87, *p*=0.006, Tukey’s post hoc tests, hM3D versus mCherry, *p*=0.010, hM3D versus hM4D, *p*=0.025; **Supplementary Figure S2D**). Additionally, hM3D mice displayed fewer entries to the open arms of the EPM compared to hM4D mice (**Figure 1J**, one-way ANOVA, *F*_2,38_=4.86, *p*=0.013, Tukey’s post hoc test, hM3D versus hM4D, p<0.011). Moreover, hM3D mice spent significantly more time in the closed arms of the EPM compared to mCherry control and hM4D mice (**Supplementary Figure S2E**). We saw no group differences in the number of entries to the closed arm (**Supplementary Figure S2F**), middle platform (*F*_2,38_=2.130, *p*=0.133; data not shown), or in the time spent in the middle platform (*F*_2,38_=1.749, *p*=0.187; data not shown). All three experimental groups showed equivalent total distance travelled in the EPM apparatus (**Figure 1K**). We found a significant effect of sex on the distance travelled in the EPM, with hM4D males travelling significantly less distance compared to hM4D females (*F*_1,35_=8.23, *p*=0.007; Sidak’s multiple comparisons test *p*>0.109; no main effect of treatment or interaction. For a discussion of sex differences, see supplemental materials). Importantly, chemogenetic activation of the IL did not affect performance in the OFT or in the EPM (**Figure 1M-R**). Taken together, our data suggest that acute activation of DP/DTT, but not IL neurons results in increased anxiety-like responses in the OFT and EPM.

### Chemogenetic activation of DP/DTT increases passive coping behavior

To examine whether chemogenetic modulation of DP/DTT activity affects passive coping behavior, we next subjected mice to the TST (**Figure 2A-B**). The hM3D mice spent significantly more time immobile compared to mCherry and hM4D groups following C21 injection (**Figure 2C-D**, one-way ANOVA, *F*_2,38_=5.07, *p*=0.11; Tukey’s post hoc tests, hM3D vs mCherry, *p*=0.041, hM3D vs hM4D, *p*=0.017; **Supplementary Figure S2G-H**), consistent with increased passive coping in the hM3D group. All three groups showed equivalent length of the longest bout of immobility, and number of transitions from mobility to immobility states (**Supplementary Figure S2I-J**). Chemogenetic activation of IL did not affect immobility in the TST (**Figure 2E-F**).

**Figure 2.**
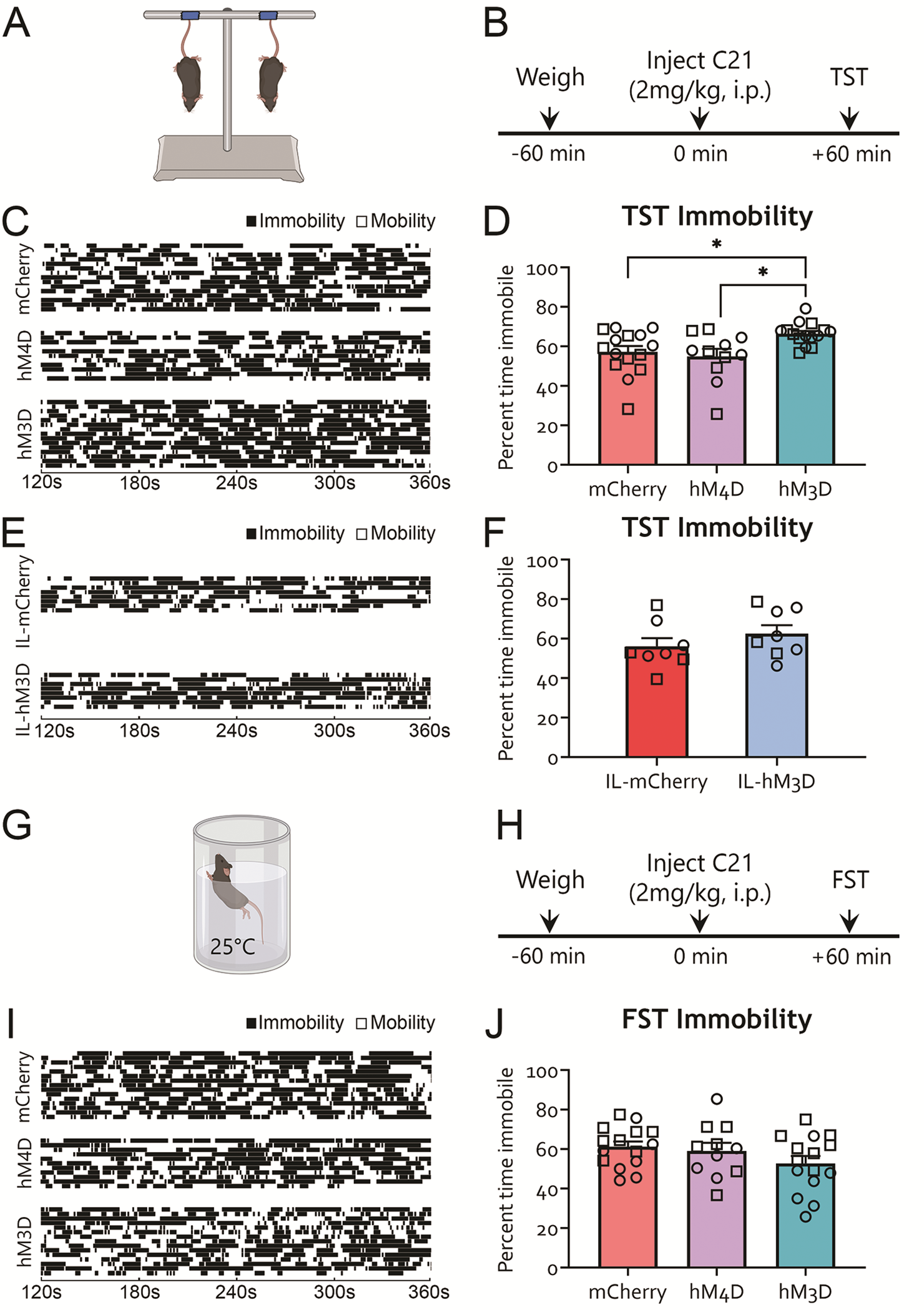
Chemogenetic activation of the DP/DTT increases immobility in the tail suspension test, but have no effect on forced swim behaviors. **A-B.** TST experimental timeline. Mice previously injected with hM3D, hM4D or control mCherry constructs in the DP/DTT were injected with the DREADD agonist C21 one hour prior to undergoing the TST. **C.** Representative raster plots for each mouse indicating periods of immobility (black) and mobility (white) over the tail suspension test duration. **D.** The average percent of time spent immobile during the TST was significantly greater in hM3D mice compared to mCherry and hM4D mice (but not different between mCherry and hM4D, p=0.838). **E-F.** Effects of IL DREADD soma manipulation on the TST. Mice previously injected with hM3D or control mCherry constructs in the IL were injected with the DREADD agonist C21 one hour prior to undergoing the TST. **E.** Representative raster plots for each mouse indicating periods of immobility (black) and mobility (white) over the TST duration. **F.** IL-hM3D mice spent the same amount of time immobile in the TST as IL-mCherry mice (*t_14_*=1.102, *p*=0.2892). **G-H.** FST experimental timeline. Mice previously injected with hM3D, hM4D or control mCherry constructs in the DP/DTT were injected with the DREADD agonist C21 one hour prior to undergoing the forced swim test. **I.** Representative raster plots for each mouse indicating periods of immobility (black) and mobility (white) over the FST duration. **J.** There was no effect of treatment on percent immobility in the forced swim test (one-way ANOVA, *F*_2,38_=1.722, p=0.192). While we found that hM3D females had a lower percent time spent immobile compared to males (see results), separation of the data by sex did not reveal group differences (One-way ANOVA, females F_2,16_= 2.816, p= 0.0896; males F_2,19_=1.042, p= 0.3722). *p<0.05. Male (square) and female (circle) individual datapoints are displayed for transparency (see text for details).

To expand our findings of increased passive coping behavior in the TST following DP/DTT activation, we conducted the FST (**Figure 2G-H**). C21 treatment did not affect time spent immobile (**Figure 2I-J**), longest bout of immobility or the number of transitions between mobility and immobility states (**Supplementary Figure 2K-N**) in any of the experimental groups in the FST. When we analyzed sex differences, we found that hM3D females spent significantly less time immobile than hM3D males and hM4D females (Two-way ANOVA, F_1,35_=6.561, *p*=0.015; Sidak’s multiple comparisons test, hM3D females versus hM3D males *p*=0.0093; hM3D females versus hM4D females *p*=0.0482). Taken together, our results demonstrate that activation of DP/DTT neurons promotes passive coping behavior in the TST, but did not alter behavior in the FST.

### Chemogenetic activation of DP/DTT activity facilitates auditory fear extinction

We next sought to explore whether manipulating DP/DTT activity might affect fear memory. To test this, we first inhibited or activated the DP/DTT one hour prior to training mice to associate an auditory tone with an electric foot shock (**Figure 3A**). During fear acquisition, chemogenetic manipulation of DP/DTT activity did not affect baseline freezing (one-way ANOVA, F_2,38_=2.367, *p*=0.107; data not shown), or the mean percent freezing across all tones (**Figure 3B**). However, we found that hM4D mice displayed a right shift in their fear learning curve, with significantly higher freezing during tone 4 and 6 of the training protocol compared to hM3D mice (**Figure 3C**, two-way rmANOVA, significant effect of tone *F*_5,190_=100.6, *p*<0.001; Tukey’s post hoc tests, minute 4 hM3D versus hM4d, *p*=0.012, and minute 6 hM3D versus hM4D, *p*=0.012) suggesting a slight facilitation of auditory fear acquisition upon DP/DTT inhibition. We found a significant main effect of sex on average freezing to tone (Two-way ANOVA, F_1,35_=7.786, *p*=0.009, see supplemental materials).

**Figure 3.**
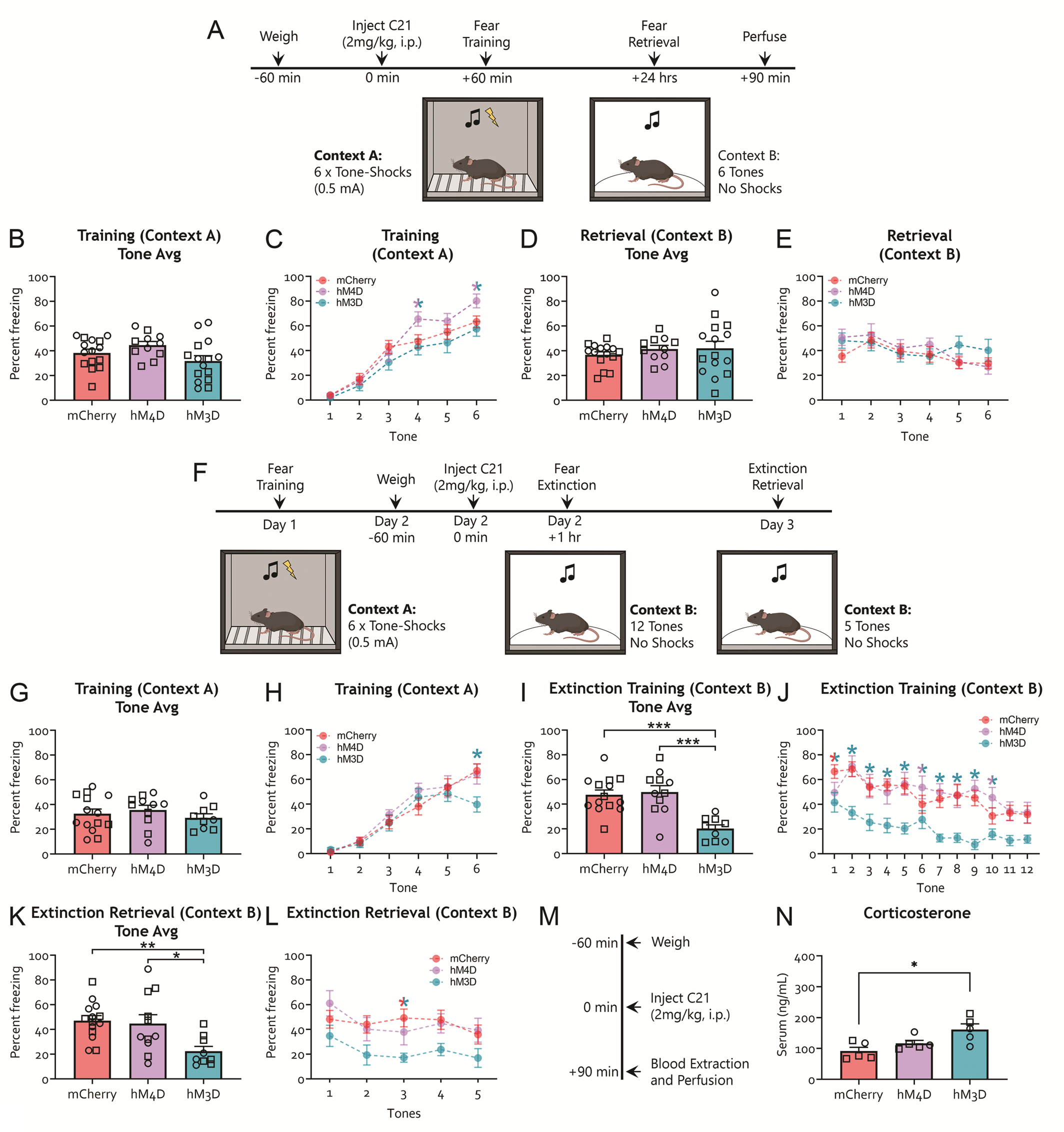
Chemogenetic inhibition of the DP/DTT has modest effect on auditory fear acquisition, but activation of the DP/DTT facilitates within-session extinction, extinction retrieval and increases serum corticosterone levels. **A-E.** DP/DTT manipulation during auditory fear acquisition. **A.** Experimental timeline. Mice previously injected with hM3D, hM4D or control mCherry constructs in the DP/DTT were injected with the DREADD agonist C21 one hour prior to undergoing auditory fear conditioning. During acquisition, mice were placed in a sound-attenuated fear conditioning chamber and received 6 tone-shock pairings (0.5mA). Twenty-four hours later, auditory fear memory was evaluated in a different context in the absence of shocks. **B.** During fear acquisition, DP/DTT manipulations did not affect average tone freezing (one-way ANOVA, F_2,38_=2.828, *p*=0.072). **C.** hM4D mice displayed significantly more freezing than hM3D mice at tone presentations 4 and 6 during acquisition. Two-way rmANOVA found no main effect of treatment *F*_2,38_=2.828, *p*=0.072, but a significant main effect of tone (*F*_5,190_=100.6, *p*<0.001, and no treatment by tone interaction: *F*_10,190_=1.633, *p*=0.998). **D-E.** During fear retrieval, DP/DTT manipulations did not affect freezing during average tone (**D**; one-way ANOVA, F_2,38_ =0.507, *p*=0.607). Individual tone analysis (**E**) showed no main effect of treatment (Two-way rmANOVA, *F*_2,38_=0.507, *p*=0.607), but a significant main effect of tone (*F*_5,190_=4.00, *p*=0.002, no treatment by tone interaction: *F*_10,190_=1.213, *p*=0.285), likely driven by within-session extinction. indicates a significant difference between hM3D and hM4D groups (*p*<0.05). **F-L.** DP/DTT manipulation during fear extinction. **F**. Experimental timeline. Mice previously injected with hM3D, hM4D or control mCherry constructs in the DP/DTT (no prior behavior) underwent fear acquisition (as previously; 6 tone-shock pairings). The following day, mice were injected with the DREADD agonist C21 one hour prior to undergoing extinction training. During extinction training, mice were placed in an alternate context (context B) and received 12 tone presentations. The following day, fear memory was evaluated during extinction retrieval, in the absence of C21. During extinction retrieval, mice were placed in context B and received 5 tone presentations. **G-H**. During auditory fear acquisition, mice from all groups displayed similar average tone (**G**; one-way ANOVA *F_2,31_* = 0.5886, *p*=0.5612) but decreased freezing in tone 6 in the hM3D group (**H**; Two-way rmANOVA significant effect of tone *F_5,155_*=74.69, *p<0.0001* and interaction *F_10,155_*=1.984, *p*=0.0385; but no main effect of treatment *F_2,31_*=0.8356, *p*=0.4431; tone 6 mCherry vs hM3D *p*=0.0023 and hM3D vs hM4D *p*=0.0109) freezing. **I-J**. During extinction training, 1 hour after C21 treatment, hM3D mice showed decreased average tone freezing (**I;** Two-way rmANOVA main effect of treatment and tone but no interaction *F_22,341_*=1.072, *p*=0.3758) and tone-by-tone (**J**) freezing compared to hM4D and mCherry control mice. **K-L**. During extinction retrieval, hM3D mice froze less compared to hM4D and mCherry control mice in the average tone (**K**) and tone-by-tone (**L;** Two-way rmANOVA significant main effect of treatment and tone but no interaction *F_8,124_*=0.6037, *p*=0.7733) analyses. Minimal sex differences were found in this dataset (see Supplementary Materials). **M**. Experimental timeline for serum corticosterone assay. Mice previously injected with hM3D, hM4D or control mCherry constructs in the DP/DTT were injected with the DREADD agonist C21 and euthanized 90 minutes later. Blood was extracted at the time of perfusion for corticosterone measurements. **N.** Serum corticosterone was significantly higher in hM3D mice compared to mCherry mice (but no change between mCherry and hM4D, *p*=0.440, hM4D vs hM3D, *p*=0.103). indicates a significant group difference between mCherry and hM3D conditions; indicates a significant group difference between hM3D and hM4D conditions; indicates a significant difference between hM3D and all other groups. *p<0.05. Male (square) and female (circle) individual datapoints are displayed for transparency.

Auditory fear memory retrieval was evaluated 24h later in a different context (**Figure 3A**). All three experimental groups showed similar baseline freezing (one-way ANOVA, *F*_2,38_=0.153, p=0.858; data not shown), and equivalent freezing to the tones (**Figure 3D-E**). Overall, these data suggest that manipulating DP/DTT activity has no major impact on auditory fear acquisition, except for a marginal facilitation of fear acquisition following DP/DTT inhibition.

To test whether DP/DTT manipulation might affect fear extinction and retrieval, we infused AAV vectors encoding either hM3D, hM4D or mCherry into the DP/DTT of a separate cohort of mice, and trained them in auditory fear conditioning as previously (**Figure 3F-H**). The next day, we injected C21 one hour prior to extinction training. During extinction training, hM3D animals froze less than hM4D and mCherry animals (**Figure 3I**: One-way ANOVA *F_2,31_*=12.91, p<0.0001, mCherry versus hM3D *p*=0.0003, hM3D versus hM4D *p*=0.0002; 3J: Two-way rmANOVA main effect of treatment *F_2,31_*=12.91, *p*<0.0001 and tone *F_11,341_*=7.869, *p*<0.0001 only; mCherry versus hM3D *p*=0.0003, hM3D versus hM4D *p*=0.0002), suggesting that DP/DTT activation impairs auditory fear retrieval and facilitates within session extinction. The next day, animals underwent extinction retrieval in the absence of C21. At extinction retrieval, hM3D animals showed less freezing than both hM4D and mCherry animals (**Figure 3K**: One-way ANOVA *F_2,31_*=5.853, p=0.0070, mCherry versus hM3D p=0.0078, hM3D versus hM4D p=0.0245; 3L: Two-way rmANOVA significant main effect of treatment *F_2,31_*=4.526, *p*=0.0189 and tone *F_4,124_*=3.021, *p*=0.0204 only; mCherry versus hM3D *p*=0.0255, hM3D versus hM4D *p*=0.0385). Overall, these data show that DP/DTT activation leads to reduced fear recall and a facilitation of fear extinction learning and extinction retrieval.

### Chemogenetic activation of DP/DTT increases serum corticosterone levels

Given the important role of the mPFC in regulating the hypothalamic-pituitary-adrenal (HPA) axis response to emotional stress [72,79–81], and the regulation of stress-induced sympathetic responses by the DP/DTT [68], we next tested the effects of DP/DTT chemogenetic modulation on serum levels of the stress hormone corticosterone. We injected C21 into a subset of mCherry, hM3D, and hM4D mice that remained in their home cage, and subsequently extracted blood to evaluate serum corticosterone (**Figure 3M**). Notably, these mice had not undergone any previous behavioral testing. Interestingly, hM3D mice displayed higher serum corticosterone levels compared to mCherry control mice (**Figure 3N**, one-way ANOVA, F_2,12_=6.325, *p*=0.013, Tukey’s post hoc tests, hM3D versus mCherry, *p*=0.011). Chemogenetic inhibition of DP/DTT activity did not affect baseline serum corticosterone levels (**Figure 3N**). These data indicate that activation of DP/DTT neurons is sufficient to trigger a neuroendocrine stress response by increasing serum corticosterone levels. We validated our chemogenetic manipulations using cFos immunohistochemistry and slice electrophysiology (**Supplementary Figure S3**).

### Neurochemical profile and mapping of DP/DTT downstream projections

To explore potential downstream effectors mediating the effects of DP/DTT activation on affective, fear behavior and neuroendocrine responses, we first analyzed brain-wide patterns of DP/DTT axonal innervation with the help of the mCherry tag in AAV-injected mice (**Figure 4A-C**). Projections from the DP/DTT were observed in the lateral septum (medial and lateral nuclei), indusium griseum (IG), BNST, dorsal endopiriform nucleus (DEn), thalamus (nucleus reuniens, RE; paraventricular nuclei, PVT; submedius nucleus, Sub), hypothalamus (DMH; lateral hypothalamus, LH; paraventricular nucleus, PVN; posterior hypothalamus, PH; and retromammillary nuclei, RM) and brainstem (periaqueductal gray, PAG) (**Figure 4A-C**). Posterior brainstem sections containing the raphe nuclei also showed DP/DTT mCherry+ projections (data not shown).

**Figure 4.**
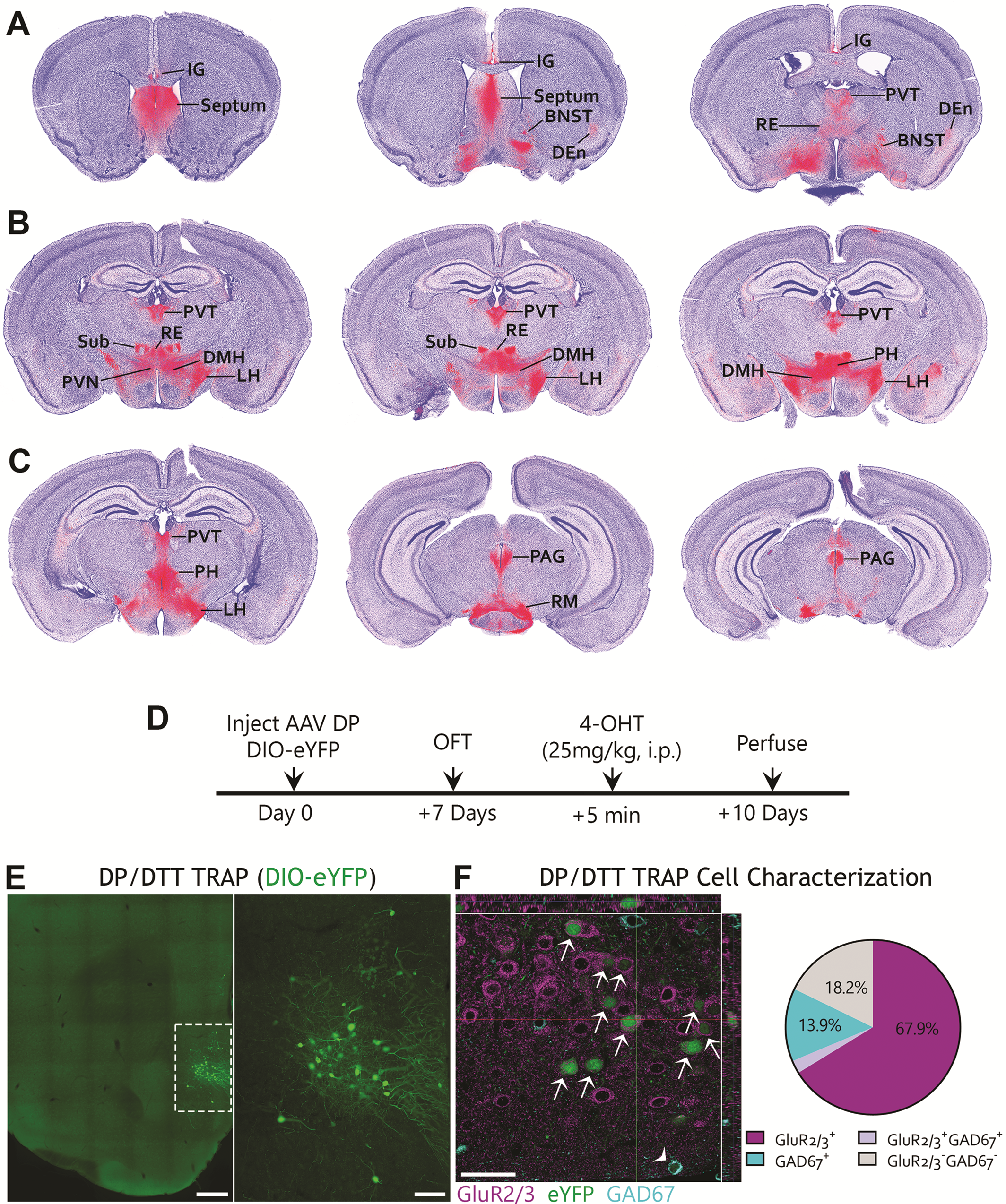
Mapping of DP/DTT downstream projections and characterization of activity-tagged DP/DTT neurons following the OFT. **A-C.** Representative projections of the DP/DTT in anterior, intermediate, and posterior sections. **A.** In anterior sections, we observed notable viral expression in the indusium griseum (IG), septum, bed nucleus of the stria terminalis (BNST), dorsal endopiriform nucleus (DEn), nucleus reuniens (RE) and paraventricular thalamus (PVT). **B.** In intermediate sections, we observed continued viral expression in the PVT and RE. Viral expression was also observed in the paraventricular nucleus (PVN) submedius thalamic nucleus (Sub), dorsomedial hypothalamus (DMH), lateral hypothalamus (LH), and posterior hypothalamus (PH). **C.** In the most posterior sections we examined, viral expression was detected in the periaqueductal gray (PAG), retromammillary nuclei (RM), PVT, LH and PH. In posterior brainstem sections, we also observed DP/DTT projections to the raphe nuclei (data not shown). **D-F**. Characterization of neurochemical profile and downstream projections patterns from activity-tagged DP/DTT neurons following the OFT. **D**. Experimental timeline. TRAP2 mice were infused with AAV expressing a DIO-YFP construct in the DP/DTT. Seven days later, animals underwent the OFT and were injected with 4HT tamoxifen immediately afterwards. Ten days later, animals were perfused and their brains processed for immunohistochemistry against GFP, glutamate receptor 2/3 (GluR2/3) and glutamate decarboxylase 67 (GAD67). **E**. Representative image of DP/DTT activity tagged neurons (green). Scale bars:400µm (left), 100µm (right inset). **F**. Left: Representative image showing DP/DTT OFT activity-tagged neurons (green) co-stained with GluR2/3 (magenta; arrows) or GAD67 (cyan; arrow heads). Scale bar:50µm. Right: Distribution of DP/DTT OFT activity-tagged neurons indicates that these cells predominantly co-label with GluR2/3 (67.9%), with 13.9% of neurons co-labeled with GAD67, 18.2% negative for either marker, and a minority of cells (2.5%) co-labeled for both GluR2/3 and GAD67 (n=6).

To determine which DP/DTT neuronal subpopulations and downstream targets are actively recruited during behavior, we next used viral tracing in Fos^2A-iCreER^ (TRAP2) transgenic mice [82,83] to tag DP/DTT neurons active during the OFT. We infused an AAV expressing Cre-dependent eYFP (AAV-DIO-eYFP) into the DP/DTT of TRAP2 mice and, following recovery, injected animals with tamoxifen immediately after OFT testing to tag the soma and axons of activated DP/DTT neurons through YFP expression (**Figure 4D-E**). We found that 67.9% of DP/DTT neurons active during OFT testing co-labeled with a glutamatergic marker, 13.9% with a GABAergic marker, and 2.5% of labeled cells were positive for both markers (**Figure 4F**). Overall, this points to OFT activating a heterogeneous but predominantly glutamatergic DP/DTT cell population.

### Optogenetic activation of the DP/DTT-DMH pathway acutely suppresses freezing but does not alter affective behavior

We next wanted to test whether one of the main downstream pathways revealed by our previous analysis, the DP/DTT-DMH pathway [68], might underlie our effects on affective and fear extinction behavior. We optogenetically targeted the DP/DTT-DMH pathway by expressing the excitatory opsin channelrhodopsin (ChR2) or control GFP into the DP/DTT and implanting an optic fiber over the DMH, and tested the animals in the OFT, EPM and auditory fear (**Figure 5A, Supplementary Figure S4A-B**). Optical activation of the DP/DTT-DMH pathway did not alter behavior in the OFT (**Figure 5B-D**) or EPM tasks (**Figure 5E-G**). In contrast, activation of the DP/DTT-DMH during tone exposure at extinction learning significantly suppressed freezing responses (**Figure 5H-I**: *t_10_*=7.119, p<0.0001; **Figure 5J**: Two- way rmANOVA significant main effect of stimulation *F_1,10_*=50.68, *p*<0.0001; tone *F_3.961,39.61_*=3.026, *p=*0.0291, and interaction *F_11,110_*=1.883, *p*=0.0491; Tone 1 *p*=0.0042, Tone 2 *p*=0.0461) without affecting extinction retrieval (**Figure 5K-L**). To test whether this suppression of freezing was specific to the tone- shock association, we fear conditioned a separate cohort of mice and activated the DP/DTT-DMH pathway in the alternate context in the absence of tones (**Figure 5M**). DP/DTT-DMH activation led to a reduction in baseline freezing behavior (**Figure 5N**; Two-way rmANOVA significant main effect of stimulation *F_1,11_*=15.88, *p=*0.0021 and interaction *F_1,11_*=9.375, *p*=0.0108 only; ChR2 *p*=0.0011) and an increase in rearing (**Figure 5O**; Two-way rmANOVA significant main effect of stimulation *F_1,11_*=16.30, *p=*0.002 and interaction *F_1,11_*=11.28, *p*=0.006 only; ChR2 *p*=0.0008). These data suggest that activation of the DP/DTT-DMH pathway acutely suppresses freezing at baseline and during extinction training without affecting extinction retrieval.

**Figure 5.**
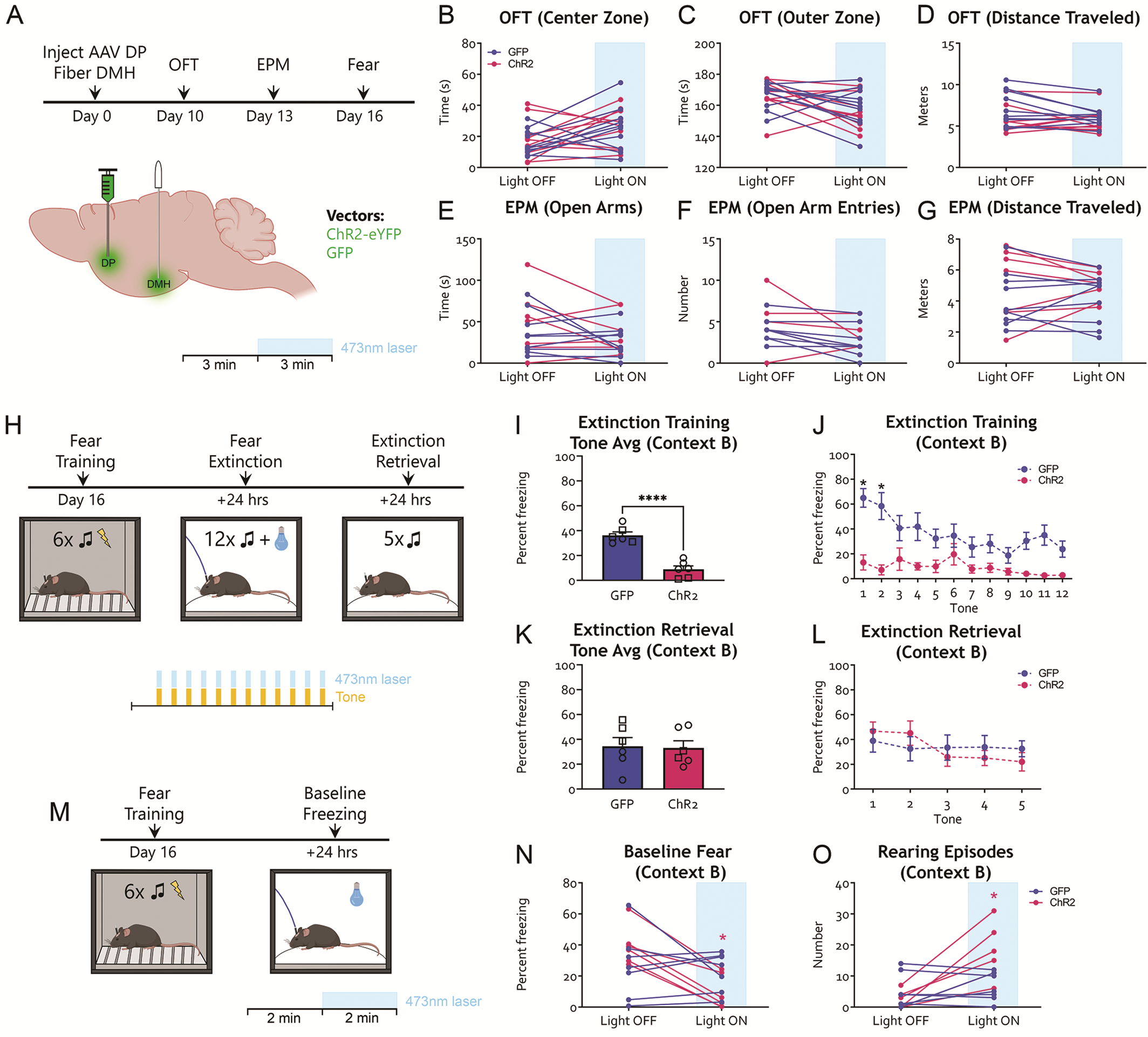
Optogenetic activation of the DP/DTT-DMH pathway acutely suppresses freezing but has no effect on affective behavior. **A**. Experimental timeline. Mice were injected with ChR2 or control GFP AAV constructs in the DP/DTT and implanted with an optic fiber in the DMH. Ten days later, animals underwent behaviour, starting with the OFT. Three days later, animals went through the EPM and, following another 3 days, one of two auditory fear conditioning protocols (see below). **B-D**. Behavior in the OFT. **B**. Light activation did not affect the time spent in the center of the open field between groups (Two-way rmANOVA significant main effect of stimulation *F_1,18_*= 5.657, *p*=0.0287; but no main effect of treatment *F_1, 18_*=0.1118, *p*=0.7420 or interaction *F_1,18_*=0.02443, *p*=0.8775; no significant posthoc tests, p>0.17). **C**. Light activation did not affect the time spent in the outer zone of the open field between groups (Two-way rmANOVA significant main effect of stimulation *F_1,18_*= 6.549, *p*=0.0197; but no main effect of treatment *F_1, 18_*=0.11240, *p*=0.7288 or interaction *F_1,18_*=0.1744, *p*=0.6811; no significant posthoc tests, p>0.097). **D**. The total distance travelled during the test was not affected by light activation (Two-way rmANOVA no main effect of treatment *F_1, 18_*=2.724, *p*=0.1162; stimulation *F_1,18_*= 3.054, *p*=0.0976; or interaction *F_1,18_*=2.728, *p*=0.1160). **E-G**. Behavior in the EPM. Light activation did not affect the time spent in the open arms of the EPM (**E**; Two-way rmANOVA no main effect of treatment *F_1,14_*=0.9766, *p*=0.3398; stimulation *F_1,14_*= 2.631, *p*=0.1271; or interaction *F_1,14_*=0.0100, *p*=0.9217), the number of entries to the open arms of the EPM (**F**; we note a non-significant trend toward a decrease in open arm entries in the EPM for the ChR2 animals: Two-way rmANOVA significant effect of stimulation *F_1,14_*= 10.53, *p*=0.0059; but no main effect of treatment *F_1,14_*=3309, *p*=0.5743 or interaction *F_1,14_*=0.1645, *p*=0.6912; Šídák’s multiple comparisons test, ChR2 p=0.057, GFP p=0.097) or the total distance travelled (**G**; Two-way rmANOVA no main effect of treatment *F_1,14_*=1.207, *p*=0.2904; stimulation *F_1,14_*= 0.0676, *p*=0.7986; or interaction *F_1,14_*=0.00511, *p*=0.0940). **H-L**. Effect of DP/DTT-DMH activation on fear extinction and extinction retrieval. **H**. Experimental timeline for the fear extinction cohort. Three days after the EPM, mice were trained in auditory fear conditioning as described previously (6 tone-shock pairings). We note a significant effect of tone on fear training in this cohort (data not shown; Two-way rmANOVA significant effect of tone *F_3.537, 35.37_*=50.62, *p<0.0001*; but no main effect of treatment *F_1,10_*=1.792, *p*=0.2103 or interaction *F_5,50_*=0.5963, *p*=0.7029). The following day, mice underwent extinction training in context B (12 tones, as previously) with light activation for the duration of each tone. On the next day, they underwent extinction retrieval in context B (5 tones, same protocol as before) in the absence of light. **I**. During extinction training, DP/DTT-DMH optogenetic activation reduced average freezing (**I**) and freezing across tones (**J**). **K-L**. During extinction retrieval in the absence of light, average tone freezing (**K**; *t_10_*=0.1392, *p*= 0.8921) and freezing across tones (**L;** Two-way rmANOVA no significant main effect of treatment *F_1,10_*=0.0193, *p*=0.8921; tone *F_2.707,27.07_*=2.575, *p=0.0798* or interaction *F_4,40_*=1.592, *p*=0.1953) were similar between groups. **M-O**. Effect of DP/DTT-DMH activation on baseline fear after conditioning. **M**. Experimental timeline for the fear baseline cohort. Three days after the EPM, a separate cohort of mice was trained in auditory fear conditioning as previously (6 tone-shock pairings). The following day, mice were placed in context B in the absence of tones, and after 2 minutes of baseline exploration the light was turned on for 2 minutes. **N**. DP/DTT-DMH optogenetic activation decreased freezing in the absence of tones. Two-way rmANOVA significant main effect of stimulation and interaction, but not virus *F_1,11_*=0.006, *p=*0.9396; Šídák’s multiple comparisons test, light OFF minus Light ON ChR2 *p*=0.0011, GFP *p*=0.7607. **O**. DP/DTT-DMH optogenetic activation increased rearing instances. Two-way rmANOVA significant main effect of stimulation and interaction, but not virus *F_1,11_*=0.8904, *p=*0.3656; Šídák’s multiple comparisons test, light OFF minus Light ON ChR2 *p*=0.0008 GFP p=0.8609. *p<0.05. Male (square) and female (circle) individual datapoints are displayed for transparency.

## Discussion

Here we examined the effects of modulating the activity of DP/DTT neurons to anxiety-, depression-like and fear behavior in stress-naïve mice. We found that chemogenetic activation of the DP/DTT reduced the time spent in the center zone of the OFT and in the open arms of the EPM. DP/DTT activation also increased passive coping in the TST, but did not affect behavior in the FST. DP/DTT activation facilitated fear extinction training and extinction retrieval, whereas DP/DTT-DMH activation acutely suppressed freezing. Interestingly, DP/DTT activation led to an increase in serum corticosterone levels, suggesting that the DP/DTT also modulates HPA axis activity. DP/DTT projection mapping showed strong innervation of lateral septum, thalamus, hypothalamus, and brainstem, consistent with tracing studies [33,67,84,85].

Our data showed that activation of the DP/DTT increased anxiety-like behavior in the OFT and EPM. A considerable body of work implicates the rodent mPFC in the regulation of anxiety-like behavior [7,13,22–29,86,14–21], with contradictory findings supporting anxiogenic [18–23,53–55] or anxiolytic [24–27] roles. Manipulations selective to the vmPFC/IL also see conflicting effects on EPM and open field performance [21,24,26,60]. Recently, Chen and colleagues found that optogenetic activation of IL soma drives anxiety-like behavior in the EPM and open field [60], similar to our data with DP/DTT activation. In contrast, they found that optical stimulation of the DP increased entries to the open arms in the EPM and time spent in the center of the open field [60], contrary to our findings. Our data support an anxiogenic role for the DP/DTT, which is in line with findings from Kataoka and colleagues showing that inhibition of the DP/DTT pathway abolishes stress-induced avoidance behavior [68], and our finding of increased serum corticosterone following DP/DTT activation. Importantly, the viral spread over the DP in the study by Chen and colleagues is more dorsal (spreading to IL, but not DTT) compared to ours, and targets glutamatergic neurons under the αCaMKII promoter [60], in contrast to the pan-neuronal synapsin and EF1a promoters used here and by Kataoka and colleagues [68]. These discrepancies favor additional cell-type and subregion specialization across the dorso-ventral axis of the vmPFC in regulating anxiety-like behavior and neuroendocrine responses.

DP/DTT activation increased immobility time in the TST, but not in the FST. Given the differential implications of animal weight on water-based (FST) and suspension (TST) task demands, it is possible that the relatively high weight of our animals at the time of testing (>30g for males) contributed to the differences in performance between these tasks. The mPFC has a well-established role in modulating depression-like behavior [87] in both stress-naïve [56–59,88,89] and stress conditions [88,90–95], including IL-specific manipulations [57,59]. Interestingly, Warden and colleagues identified mPFC neuronal population coding for FST active and inactive behavioral states, which reflected specialization within mPFC downstream projections to the dorsal raphe nucleus and lateral habenula [96]. Accordingly, somatic activation of the mPFC as a whole did not affect behavior in the FST [96], suggesting that further probing of specific DP/DTT projections could lead to modulation of FST behavior despite our negative findings.

Brain-wide mapping of DP/DTT projections showed strong downstream connectivity with the hypothalamus, thalamus, lateral septum and brainstem. Importantly, many of the upstream brain regions projecting to the DP [68] also regulate anxiety-like behavior in rodents, including the mediodorsal (MD) [97–100] and paraventricular (PVT) [101–105] thalamus, insular cortex (IC) [105–110], nucleus reuniens [111] and amygdala [14,112–116]. While we found that the majority of OFT activity-tagged DP/DTT neurons is glutamatergic, the precise patterns of afferent and efferent innervation of DP/DTT neuronal subpopulations and their modulation of anxiety-like behavior remains unknown.

Our data showed minimal effects of DP/DTT soma manipulation on auditory fear acquisition, but a marked reduction in freezing when the DP/DTT was activated during fear extinction, which persisted during extinction retrieval in the absence of DP/DTT manipulation. This points to the DP/DTT as sharing a similar function to IL [61,129–132, but see 133] in the modulation of fear inhibition. In contrast, a recent study elegantly showed that activation and inhibition of DP neurons tagged during auditory fear conditioning increases and reduces freezing at retrieval, respectively [122]. The opposing direction of these findings in relation to ours could be driven by a differential contribution of the whole DP/DTT (our manipulations) compared to the selective activation of DP neurons responding to fear acquisition [122], whose connectivity is presently unknown. This is also underscored by a lack of activity signatures by these fear-trapped DP neurons during extinction [122], whereas we saw an effect of DP/DTT soma manipulation on extinction learning and retrieval.

We found that DP/DTT-DMH stimulation had no effect on affective behavior in the OFT or EPM, but acutely suppressed freezing. Notably, in contrast to our DP/DTT soma manipulation, DP/DTT-DMH- mediated suppression of freezing at extinction learning did not affect extinction retrieval, suggesting that the DP/DTT effects on extinction retrieval might be mediated by a distinct downstream target. Furthermore, the increased rearing seen in the DP/DTT-DMH ChR2 animals at baseline post-conditioning might also signal stress-related behavior [123], consistent with this pathway’s role in mediating sympathetic responses and social avoidance following social defeat stress in rats [68]. Alternatively, DP/DTT-DMH activation after foot shock might be triggering flight-like behavior as recently reported for the DP-central amygdala (CeA) pathway [124]. In this comprehensive study, inhibition of the DP-CeA pathway increased freezing during fear conditioning [124], similar to our trends with DP soma inhibition. It is also important to note that the small magnitude of our fear acquisition findings might have been affected by previous exposure to stressful manipulations such as the FST. Interestingly, stimulation of the DP-CeA pathway in a safe context after conditioning reduces freezing [124], similar to our results with DP/DTT and DP/DTT-DMH stimulation. Importantly, the authors reported an anxiolytic effect of inhibition of the DP-CeA pathway in the OFT and EPM, but no effect of activation [124], suggesting that our effects of DP soma manipulation on affective behavior might be independent of the CeA. This is in line with the absence of a strong projection to CeA with our viral strategy. Finally, Chen and colleagues demonstrated that IL downstream projections to the lateral septum and central amygdala (CeA) promote and inhibit anxiety-like behavior, respectively [60], suggesting a high degree of specialization within the vmPFC for the modulation of anxiety-like behavior.

We found that chemogenetic activation of the DP/DTT was sufficient to trigger an increase in serum corticosterone levels. This is consistent with the finding that optogenetic stimulation of the DP/DTT-DMH pathway in stress-naïve animals drives sympathetic responses mimicking those triggered by social defeat stress [68], suggesting that the DP/DTT might transduce stress responses from upstream corticolimbic regions toward downstream autonomic, behavioral/motor [68] and endocrine responses. Importantly, while most of the literature supports an inhibitory effect of the mPFC over HPA secretory responses following stress [71,125–128] (but see [129]), Radley and colleagues saw a dichotomy between the dorsal and ventral mPFC in modulating stress-induced HPA activation [72]. Specifically, while dmPFC lesions increased corticosterone levels following acute restraint stress, vmPFC lesions decreased corticosterone levels [72]. This suggests that mPFC inhibition of the HPA axis response is mostly driven by the dorsal mPFC, with the ventral portion of the mPFC (possibly spanning the DP/DTT) exerting the opposite effect, consistent with our findings. Importantly, Kataoka and colleagues saw no evidence of DP/DTT inhibition affecting basal autonomic homeostasis [68], consistent with our lack of effects of DP/DTT inhibition in stress-naïve animals. Similarly, inhibition of the ventral portion of the mPFC did not affect baseline corticosterone levels [72].

While we observed that the DP/DTT can modulate neuroendocrine responses, the downstream targets mediating this response are unclear. The hypothalamus is necessary for the effects of frontal cortex stimulation on HPA activity [130], and mediates the autonomic effects of DP/DTT activation [68]. It is hypothesized that (dorsal) mPFC inhibition of HPA activity occurs through stimulation of GABAergic projections from DMH to the paraventricular nucleus of the hypothalamus (PVN) [131]. Consistent with what has been described for the vmPFC [126], we saw DP/DTT projections to hypothalamic nuclei such as the PVN, DMH and lateral hypothalamus (LH). While Radley and colleagues proposed that the stimulating effects of ventral mPFC on HPA activity might be mediated through connections with BNST [72], studies suggest an inhibitory effect of the BNST over HPA activity following acute stress [132], and fail to see recruitment of mPFC-BNST projections following stress [128,133]. Altough we see DP/DTT projections to the BNST, their contribution to the regulation of DP/DTT effects on HPA secretory activity is unknown.

Overall, our data expand an emerging literature examining the contributions of the DP/DTT to the modulation of psychosocial stress and sympathetic responses by revealing the DP/DTT as a regulator of anxiety-like, passive coping and fear extinction behaviors. As its topography allows for the integration of higher order information toward autonomic and sympathetic effectors, whose dysfunction is often a feature of anxiety disorders, the DP/DTT may be uniquely placed to mediate the effects of stress on affective function. Our findings support further specialization across the dorsoventral axis of the mPFC in the processing of anxiety- and depression-like behavior and stress responsivity, in line with rising evidence of pathway-specific modulation of anxiety-like behavior among mPFC subregions.

## Acknowledgements

We thank Hathairat Chanphao, Tejnarine Persaud, Bebhinn Treanor, and Christina Guzzo for their generous and fundamental assistance with the corticosterone ELISA experiment and use of the Guzzo lab plate reader. We would also like to thank Unza Mumtaz and Mehreen Inayat for their help with animal colony management. Some figure diagrams were created with the assistance of BioRender.com.

## Author Contributions

*Designed Research:* JJB, AK, MAC.

*Performed Research:* JJB, AK, RA, JW, FV, HP, ACS, AEC, AP, SDZ, MAC

*Analyzed/Interpreted Data:* JJB, AK, JW, FV, AEC, AP, MAC.

*Wrote the paper:* JJB, AK, MAC.

All authors reviewed and approved the manuscript.

## Competing Interests

The authors have nothing to disclose.

**Supplementary Figure S1.**
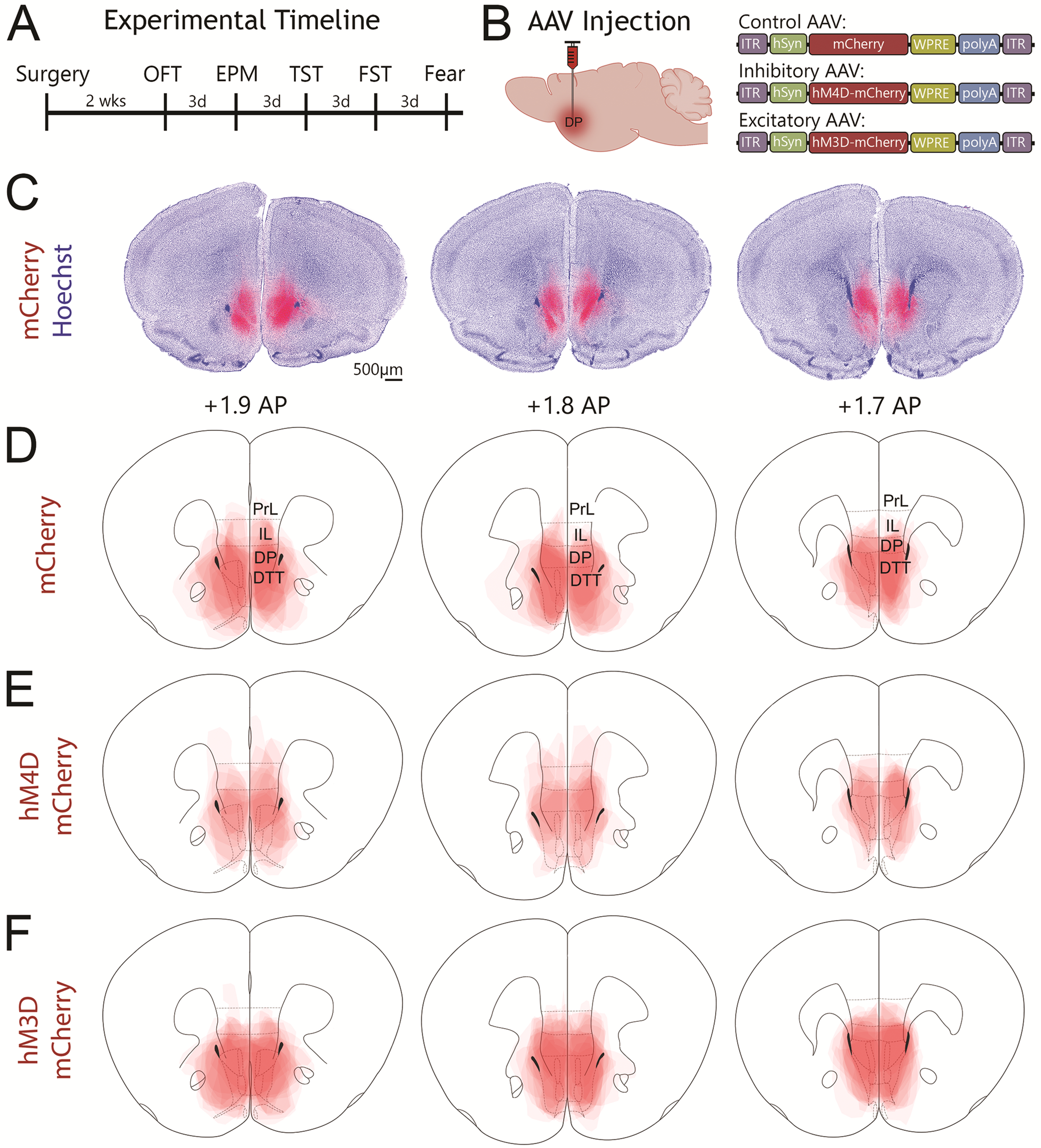
Viral expression in the DP/DTT. **A-B.** Experimental timeline. C57BL/6J mice were injected with AAV5-hSyn-mCherry, AAV5-hSyn-hM3D- mCherry or AAV5-hSyn-hM4D-mCherry into the DP/DTT. Following 2 weeks, mice underwent a battery of behavioral tests comprised of the open field test (OFT), elevated plus maze (EPM), tail suspension test (TST), forced swim test (FST) and auditory fear conditioning (Fear) with a washout period of three days between each test. **C.** Representative images of an mCherry control virus-infused brain stained for mCherry (red) and the nuclear dye Hoechst (blue). **D-F.** Diagrams depicting overlaid viral spread for all mCherry (**D**), hM4D (**E**) and hM3D-injected (**F**) mice. Scale bar = 500µm.

**Supplementary Figure S2.**
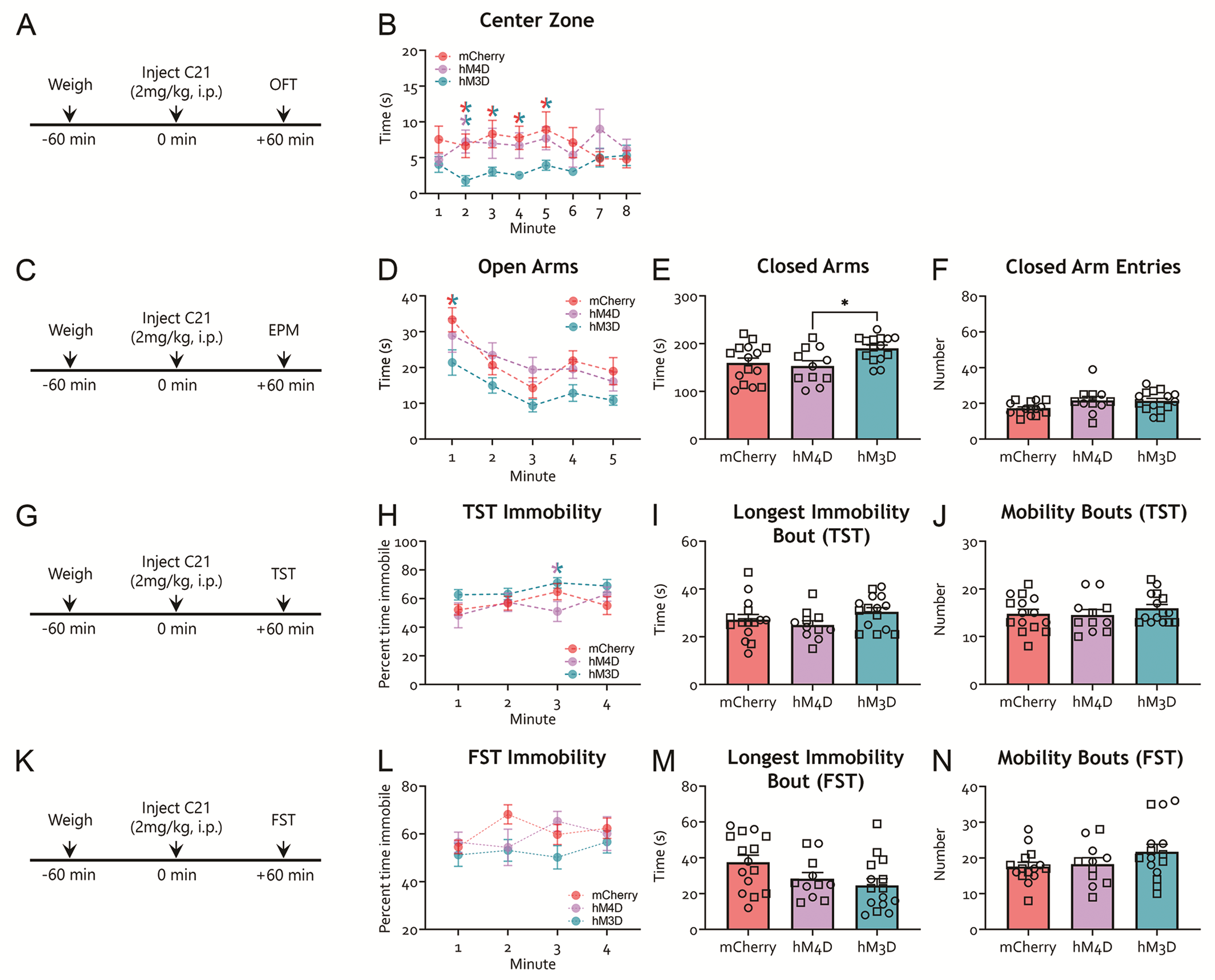
Additional behavioral metrics for DP/DTT soma manipulations on affective behavior. **A.** Experimental timeline for the OFT. Mice previously injected with hM3D, hM4D or control mCherry constructs in the DP/DTT were injected with the DREADD agonist C21 one hour prior to being placed in an open field. The time spent in the center zone (red) and the outer zone (white) was measured. **B.** Time spent in the center of the open field arena was significantly reduced in hM3D mice compared to mCherry mice during minutes 2, 3, 4, and 5. The hM3D mice also spent less time in the center of the arena compared to hM4D mice during minute 2 (two way-rmANOVA, *F*_2,38_=7.795, *p*=0.002, Tukey’s post hoc tests: minute 2, hM3D mice versus mCherry mice, *p*=0.047 and hM3D versus hM4D mice *p*=0.039; minute 3, 4 and 5, hM3D versus mCherry, *p=*0.03, 0.031 and 0.04 respectively). indicates a significant difference between hM3D and mCherry groups; indicates a significant difference between hM3D and hM4D groups. **C**. Experimental timeline for the EPM test. Mice previously injected with hM3D, hM4D or control mCherry constructs in the DP/DTT were injected with the DREADD agonist C21 one hour prior to being placed in an elevated plus maze. **D**. hM3D mice spent significantly less time in the open arms of the elevated plus maze compared to mCherry mice (minute 1) (two-way repeated measures ANOVA (rmANOVA), main effect of treatment *F*_2,38_=5.867, *p*=0.006 and minute *F*_4,152_=13.45, *p*<0.001 but no interaction *F*_8,152_=0.711, *p*=0.681; Tukey’s post hoc tests: minute 1, hM3D versus mCherry, *p*=0.008; minute 3. **E**. The amount of time spent in the closed arms of the elevated plus maze was significantly greater in the hM3D mice compared to mCherry and hM4D mice (one-way ANOVA, *F*_2,38_=4.48, *p*<0.018, Tukey’s post hoc tests, hM3D versus mCherry, *p*=0.052, hM3D versus hM4D, *p*=0.028, no difference between the mCherry control and hM4D groups: *p*=0.89). **F**. The number of closed arm entries during the test did not differ between groups (*F*_2,38_=2.730, *p*=0.078). indicates a significant group difference between mCherry and hM3D conditions. **G**. TST experimental timeline. Mice previously injected with hM3D, hM4D or control mCherry constructs in the DP/DTT were injected with the DREADD agonist C21 one hour prior to undergoing the TST. **H**. Minute-by-minute analysis of immobility on the TST found that hM3D mice spent significantly more time immobile than the hM4d mice during the third minute of the test. A two-way repeated measures ANOVA found a significant main effect of treatment (*F*_2,38_=5.077, *p*=0.011), but no main effect of minute (*F*_3,114_=1.553, *p*=0.205) or treatment by minute interaction (*F*_6,114_=0.829, *p*=0.555). Tukey’s post hoc test minute 3: hM3D mice versus hM4D mice, *p*=0.0275; indicates a significant difference between hM3D and hM4D groups. **I.** The longest continuous period of immobility did not differ between groups (one-way ANOVA *F*_2, 38_=1.874, *p*=0.167). **J.** The number of mobility-immobility transitions (mobility bouts) did not differ between groups (one-way ANOVA *F*_2,38_=0.614, *p*=0.547). **K**. FST experimental timeline. Mice previously injected with hM3D, hM4D or control mCherry constructs in the DP/DTT were injected with the DREADD agonist C21 one hour prior to undergoing the FST. **L.** Minute-by-minute analysis of immobility on the FST. Percent immobility did not differ between groups (Two-way rmANOVA, no main effect of treatment on percent immobility *F*_2,38_=1.723, *p*=0.192, no main effect of minute *F*_3,114_=1.266, *p*=0.289, and no treatment by minute interaction *F*_6,114_=1.485, *p*=0.189). **M.** We noted a non-significant trend of treatment effect on the longest bout of immobility (one-way ANOVA, *F*_2,38_=3.22, p=0.051). **N.** The number of mobility-immobility transitions (mobility bouts) did not differ between groups (one-way ANOVA, *F*_2,38_=1.722, p=0.192). *p<0.05. Male (square) and female (circle) individual datapoints are displayed for transparency (see text for details).

**Supplementary Figure S3.**
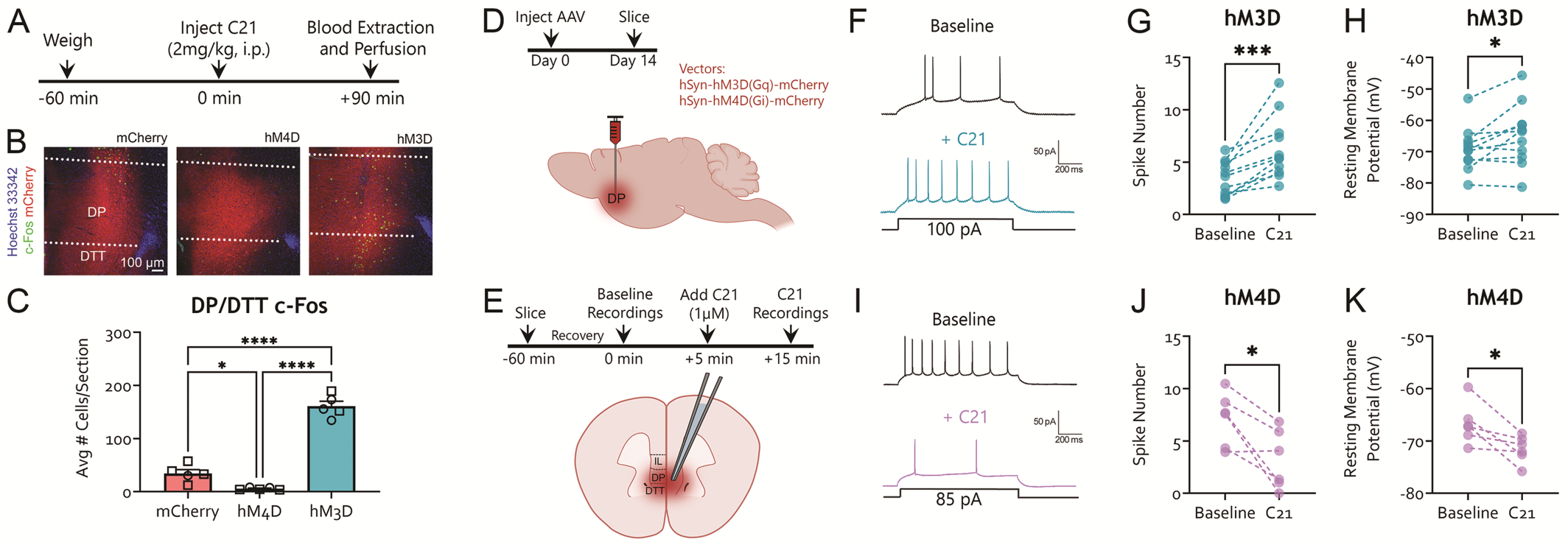
Confirmation of C21 effects on DP/DTT pyramidal neurons. **A-C.** Validation of DP/DTT soma manipulations using cFos immunohistochemistry. **A.** Experimental timeline for cFos analysis. Mice previously injected with hM3D, hM4D or control mCherry constructs in the DP/DTT were injected with the DREADD agonist C21 and euthanized 90 minutes later. **B**. Representative photomicrographs showing c-Fos (green) and mCherry (red) expression in the DP/DTT following C21 treatment. Note the significant number of c-Fos^+^ neurons in the hM3D mice. Hoechst 33342 (blue) was used as a nuclear marker. The lateral ventricle (dense blue stain on the right of each photomicrograph) was used as a landmark for the DP/DTT. **C.** hM3D mice had significantly more c-Fos+ neurons in the DP/DTT than mCherry and hM4D mice (one-way ANOVA, *F*_2,12_=143.9, *p*<0.001, Tukey’s post hoc tests, hM3D versus mCherry, *p*<0.001). Furthermore, mCherry mice also had significantly more c-Fos+ neurons in the DP than hM4D mice (hM4D versus mCherry, *p*=0.030, hM3D versus hM4D, *p*<0.001). Scale bar: 100µm. Male (square) and female (circle) individual datapoints are displayed for transparency. **D-K**. Validation of DP/DTT soma manipulations using slice electrophysiology. **D**. Experimental design. AAV constructs encoding hM3D-mCherry or hM4D-mCherry were injected into the DP and slices were made at least 14 days later. **E.** Experimental timeline for electrophysiological recordings. Slices were made and allowed to recover for 60 minutes prior to collecting baseline recordings in mCherry^+^ neurons. After a stable baseline recording of depolarization-induced firing, C21 (1 µM) was bath applied for at least 10 minutes and the last 5 minutes of the recording period were compared to the baseline. **F.** Representative traces of action potentials recorded from an hM3D^+^ neuron at baseline and after application of C21. **G.** The number of action potentials evoked by somatic current injection (100pA) in hM3D^+^ transfected neurons was significantly increased following bath application of C21 (paired t-test, t_10_=4.604, p=0.001; n=11 cells, 4 mice). **H.** The resting membrane potential (RMP) of hM3D^+^ neurons became significantly more depolarized following bath application of C21 (paired t-test, t_10_=2.77, p=0.02). **I.** Representative traces of action potentials recorded from an hM4D^+^ neuron at baseline and after application of C21. **J.** The number of action potentials evoked by somatic current injection (85pA) in hM4D^+^ neurons was significantly decreased following bath application of C21 (paired t-test, t_5_=3.42, p=0.019; n=6 cells, 3 mice). **K.** The RMP of hM4D^+^ neurons became significantly more hyperpolarized following bath application of C21 (paired t-test, t_5_=3.24, p=0.023). *p<0.05, ***p<0.001

**Supplementary Figure S4.**
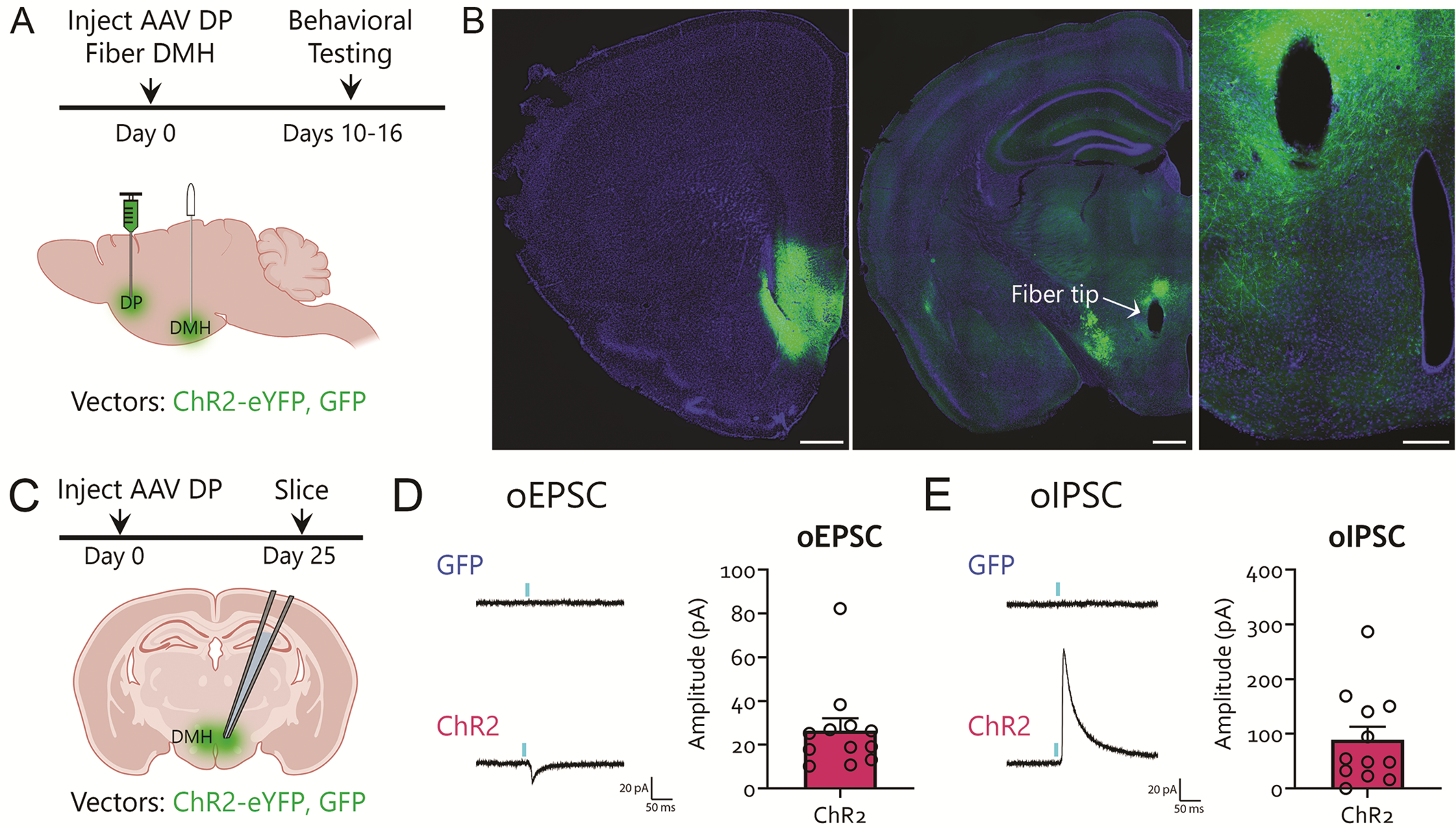
Validation of optogenetic manipulation of the DP/DTT-DMH pathway. **A.** Experimental design. AAV constructs encoding GFP or ChR2 were injected into the DP/DTT and an optical fiber was placed in the DMH. **B.** Representative image showing ChR2-EYFP expression in the DP (left, scale bar=500µm) and the DP axonal projections to the DMH where the optical fiber was placed (mid, scale bar=500µm). A higher-power inset of the DP/DTT-DMH projections and fiber location are shown on the right (scale bar=200µm). **C.** Slices were prepared at least one week after behavior was completed. **D-E**. Validation of DP/DTT optic activation in the DMH using slice electrophysiology. A light pulse (5ms) triggers monosynaptic excitatory and inhibitory responses in patched DMH neurons of ChR2 mice (bottom), but not GFP mice (top). While Kataoka et al. demonstrated that their DP/DTT-DMH effects were glutamatergic [68], future work is needed to determine the contribution of DP/DTT-triggered excitation and inhibition in the DMH to our reported effects.

## Supplemental Methods

### Animals

Adult male and female C57BL/6J and Fos^2A-iCreER^ (TRAP2) mice (2-4 months old; Jackson laboratory) were maintained on a 12 hour light-dark cycle (lights on at 7:00 A.M.), with experimental procedures performed during the light phase. Mice were bred in-house and weaned on postnatal day 21 with same- sex siblings (2-4 per cage). Food and water were available *ad libitum*. All experimental procedures were approved by the Animal Care Committee at the University of Toronto.

### Stereotaxic surgery and viral injections

Mice were injected with a cocktail of ketamine (100mg/kg) and xylazine (5mg/kg) to induce anesthesia. Once mice were fully anesthetized, the head was shaved and swabbed with iodine followed by 70% ethanol. Tear gel (Alcon) was applied to each eye to prevent dehydration. Mice were then secured in a rodent stereotaxic apparatus (Stoelting) using ear bars. Body temperature was maintained at 37°C throughout surgery with a far infrared homeothermic warming pad with a temperature feedback module (RightTemp® Jr, Kent Scientific). A scalpel was used to make an incision down the midline of the scalp, the connective tissue was excised, and the skull was cleaned with sterile phosphate buffered saline (PBS, pH=7.4). An autoclaved cotton tip applicator was briefly submerged in 30% hydrogen peroxide and then gently applied to the skull to identify bregma. A cordless micro drill (#58610, Stoelting) was used to make craniotomies over the DP/DTT (+1.85 anterior-posterior, ±0.30 medial-lateral, -3.52mm dorso- ventral, in reference to bregma) or IL (+1.9 anterior-posterior, ±0.26 medial-lateral, -2.9 dorso-ventral, in reference to bregma). Virus was delivered with a 500nL Neuros Syringe (#65457-02, Hamilton Company) attached to a probe holder on the stereotaxic apparatus (#751873, Harvard Apparatus). A total of 40- 100nL of virus was injected at a rate of 20nL per minute. The needle stayed in place for 5 minutes after each injection to allow for diffusion of the virus before being slowly removed from the brain. We used the following adeno-associated viruses (AAVs): AAV5-hSyn-mCherry (2.3 x 10^13^ vg/mL, Addgene #114472), AAV5-hSyn-hM4D(Gi)-mCherry (8.6 x 10^12^ vg/mL, Addgene #50475), and AAV5-hSyn-hM3D(Gq)-mCherry (1.9 x 10^13^ vg/mL, Addgene #50474), AAV1-CaMKIIa-hChR2(H134R)-EYFP (2.3 x 10^13^, Addgene #26969), AAV1-CAG-GFP (1.2 x 10^13^ vg/mL, Addgene #37825) and AAV5-Ef1α-DIO-EYFP (4.0 x 10^12^ vg/mL, UNC vector core). For optogenetic experiments, an optic fiber was implanted over the DMH (-1.62 anterior-posterior, ±0.30 medial-lateral, -5.00 mm dorso-ventral, in reference to bregma). Ketoprofen (1 mg/kg, s.c.) was used to reduce discomfort. Upon completing the viral injections, the skull was cleaned with sterile PBS and the scalp was sutured with Vetbond tissue adhesive (3M). For optogenetic experiments, the optic fiber was anchored by bone screws (HRS Scientific) and secured to the skull using Metabond cement (Parkell, cat#S380). Warmed physiological saline (0.5 to 0.7mL, s.c.) was injected at the end of each surgery to support hydration. Mice were placed in a clean cage located on a heating pad for post-surgical recovery and returned to the colony room once fully ambulatory. Ketoprofen (1 mg/kg, s.c.) was injected 24 and 48hr intervals after surgery to reduce post-surgical discomfort.

### Tamoxifen injections

4-hydroxytamoxifen (4-OHT; Cat# H4278, Sigma Aldrich) was mixed with ethanol (Cat# 6590-16, Fisher Scientific) and Cremaphor (Cat# C5135, Sigma Aldrich) to make a 10mg/mL stock solution, which was diluted 1:1 in 0.1M PBS and injected i.p. at a dose of 25mg/kg. TRAP2 mice were injected with AAV5- Ef1α-DIO-EYFP in the DP/DTT and one week later underwent OFT. Immediately after the OFT, mice were injected with 4-OHT and returned to the home cage. Animals were perfused 10 days later and their brains processed for immunohistochemistry.

### Behavioral testing

For experiments spanning Figures 1, 2 and 3A-E, following a 2-week recovery period after surgery, cohorts of mice underwent the following behavioral tests interspersed by a 3-day washout period: 1) open field test (OFT), 2) elevated plus maze (EPM), 3) tail suspension test (TST), 4) forced swim test (FST) and 5) auditory fear conditioning. Behavioral task order was defined by degree of stress, starting from the least to greater stress-inducing tasks to minimize carryover effects across tasks [134].

Correlation analyses between individual behavior in the OFT and EPM and subsequent freezing levels during fear training or retrieval in hM3D and hM4D animals showed no significant differences (p>0.18, data not shown), with the exception of a significant inverse correlation between the time spent in the center zone of the open field and fear memory retrieval in the hM4D group (R^2^=0.49, p=0.01). In these experiments, the hM3D group showed a higher number of animals (7/15) with very weak spread to the NAc compared to the hM4D group (2/11). Importantly, hM3D animals with or without NAc spread showed no significant difference in behavior in the OFT, EPM, TST or auditory fear (p>0.33, data not shown), suggesting this is not a major contributor to our effects. Five cohorts of mice were used in these experiments, and we observed consistent effects of DP/DTT manipulation across all cohorts.

For experiments in Figure 3F-L, animals underwent fear conditioning following recovery from surgery without prior behavior. Six cohorts of mice were used in these experiments, and we observed consistent effects of DP/DTT manipulation across all cohorts. In the experiments featured in Figure 5, all cohorts underwent the following behavioral tests interspersed by a 3-day washout period: 1) OFT, 2) EPM, 3) auditory fear training, followed the next day by either (a) fear extinction training and extinction retrieval (Figure 5H-L) or (b) baseline freezing testing (Figure 5M-O; see details below). Three cohorts of mice were used in each of these experimental designs, and we observed consistent effects across all cohorts.

The DREADD agonist compound 21 (C21; 2mg/kg, 0.5mg/mL, i.p.) was injected 1hr before the start of each behavioral test. Mice remained in their homecage until behavioral testing began. Detailed methods for each behavioral test are described below. Upon completing behavioral testing, mice were returned to their homecage and left undisturbed until the next test. All behavioral testing apparatuses were cleaned with 70% ethanol followed by ddH_2_0 between subjects. Behavior was automatically scored using ANY- maze software (Stoelting) (OFT, EPM), Depression Behavior Scorer (DBscorer) [135] (TST, FST) or VideoFreeze [136,137] (Med Associates) (fear).

#### Open Field Test

Exploratory and anxiety-like behaviors were measured in the OFT as previously described [138–141] with minor modifications. Mice were placed in the corner of a brightly illuminated open field arena (30cm length x 30cm width x 30cm height) and an overhead monochrome video camera (DMK 22AUC03, The Imaging Source) recorded behavior over the 8-minute test duration with ANY-maze software (version 7.1, Stoelting). The primary measures of interest included the time spent in the center zone of the maze (10cm x 10cm), time spent in the outer zone, and the total distance travelled during the test. Track plots and heat maps were generated using ANY-maze software. Optogenetic stimulation of the DP/DTT-DMH pathway (473 nm, 20 Hz, 5ms pulses) occurred after 3 minutes of baseline activity with the light off, and lasted for 3 minutes.

#### Elevated Plus Maze

Exploratory and anxiety-like behaviors were evaluated in the EPM as previously described [141,142] with minor modifications. The EPM apparatus was comprised of two open and two closed arms of identical dimensions (35cm length x 10cm width) raised 51.5cm above the floor. The closed arms had 30cm tall walls, whereas the open arms had a 0.1cm raised ledge to prevent mice from falling off the apparatus during testing. At the start of each test, mice were placed in the middle of the platform (10cm length x 10cm width) with their head facing towards one of the closed arms. A Logitech C920 1080P webcam was placed directly above the EPM and recorded behavior over the 5-minute test duration via Logi Capture Software (v 2.06.12, Logitech). Video recordings were loaded in ANY-maze software and EPM behavior was analyzed offline. The primary measures of interest included the time mice spent in the open and closed arms of the EPM and the time spent in the middle platform. The software also tracked the number of entries into each zone and the total distance travelled during the test. Track plots and heatmaps were generated using ANY-maze software. Optogenetic stimulation of the DP/DTT-DMH pathway (473 nm, 20 Hz, 5ms pulses) occurred after 3 minutes of baseline activity with the light off, and lasted for 3 minutes.

#### Tail Suspension Test

The TST was used to measure passive coping behavior as previously described [143,144]. Mice were suspended approximately 45 cm above the floor by their tails for 6 minutes. Laboratory tape was placed <1cm from the tip of each tail and secured to a horizontal stainless-steel rod attached to a laboratory retort stand. Pairs of mice were positioned approximately 40cm apart from each other during testing. A Logitech C920 1080P webcam was placed horizontally in front of the mice and recorded behavior over the test duration. The final 4 minutes of the TST was analyzed using the open-source software DBscorer, which performs automated and unbiased behavioral scoring [135]. This software converts each video frame to a binary image and compares the percent area difference between each frame, which are averaged for every second of the test duration. When the percent area difference is below a user-defined threshold value, the animal is considered immobile. Pilot experiments found that a 0.6% area threshold value was consistent with manually scored videos. The data was automatically scored in 60s time bins and consists of the percent time spent immobile, the longest continuous bout of immobility during the test (in seconds), and the number of mobility-immobility transitions. The software also automatically generated raster plots of immobility over the test duration for each mouse.

#### Forced Swim Test

The FST was used to measure depression-like behavior [145–147]. Mice were individually placed in a 1L beaker (14.5cm tall) that was filled with approximately 750mL of warmed tap water (25°C). A Logitech C920 1080P webcam was placed horizontally in front of the beaker and recorded behavior for 6 minutes. The final 4 minutes of the FST was analyzed [148,149] using DBscorer with an area threshold value of 1.0% [135]. Videos were scored in 60s time bins and measured the percent of time spent immobile, the longest continuous bout of immobility (in seconds), and the number of mobility-immobility transitions. Raster plots were also generated to indicate periods of mobility and immobility throughout the 4-minute analysis period. Upon completing the FST, mice were dried off with a towel, placed in a fresh cage on a heating blanket for 30 minutes, and returned to the colony room. The water in each beaker was changed between subjects.

#### Auditory Fear Conditioning

Auditory fear conditioning was used to evaluate whether manipulations of the DP/DTT influenced cognition. Auditory fear training occurred in a stainless-steel fear conditioning apparatus (32cm wide, 25.5cm high, 25.5cm deep) that contained shock-grid floors (36 rods, 2mm diameter; Context A). The fear conditioning apparatus was located within a sound-attenuated chamber (63.5cm wide, 36.8cm, high, 74.9cm deep; catalog #NIR-022MD, Med Associates). On the training day, a two-minute acclimation period was provided to assess baseline freezing behavior. A total of 6 auditory tones (2 kHz, 80 dB, 25s) were presented with a 60s interval and co-terminated with a 2s foot shock (0.5mA), adapted from our previous studies [64,137]. Upon completing the training session, mice were placed in their home cage and returned to the colony room until the following day.

Fear memory was evaluated 24 hours after the training session. The fear conditioning apparatus was modified for memory retrieval by installing a curved white plastic insert along the walls and a white plastic insert over the shock-grid floors (Context B). Baseline freezing behavior was evaluated in the first two minutes of the test. A total of 6 auditory tones (same parameters as training) were presented every minute to evaluate fear memory. Mice were returned to their home cage upon completing the fear memory test.

For experiments manipulating DP/DTT activity during extinction, separate cohorts of animals underwent surgery and ten days later were trained in auditory fear training as described above (no prior behavioral testing). Upon completing the training session, mice were placed in their home cage and returned to the colony room. The following day, for extinction training, animals were placed in context B and after a two- minute acclimation period were exposed to 12 tones (2 kHz, 80 dB, 25s) with a 35s interval. Mice were returned to their home cage and the following day underwent extinction retrieval. At extinction retrieval, animals were placed in context B and, and after a two-minute acclimation period, were exposed to 5 tones (2 kHz, 80 dB, 25s) with a 35s interval. For all DP/DTT soma extinction experiments, animals were perfused for histological confirmation of targeting, and all animals displaying viral spread to IL were excluded from the dataset.

For the experiments examining the effect of optogenetic stimulation of the DP/DTT-DMH pathway during <u>extinction training</u>, animals previously trained in OFT and EPM underwent fear training as described previously. The next day, during extinction training (same protocol as above), light stimulation (473 nm, 20 Hz, 5ms pulses) occurred for the duration of each of the 12 tones. Mice were returned to their home cage and the following day underwent extinction retrieval (same parameters as above) in the absence of light stimulation.

For the experiment looking at the effect of DP/DTT-DMH stimulation on <u>baseline freezing</u>, a separate cohort of animals previously trained on OFT and EPM (but not fear) underwent fear conditioning as described previously in the absence of light stimulation. The following day, animals were placed in context B and, after 2 minutes with the light off, underwent light stimulation (473 nm, 20 Hz, 5ms pulses) for 2 minutes in the absence of tones. Rearing was hand-scored by an experimenter blind to experimental conditions, whereby a rearing instance was counted when a mouse elevated both front paws off the floor (supported and unsupported rearing [123] were conflated in our analysis). In this cohort, some animals showed spread into the LS, but an excluded animal in which only the LS was targeted did not show the same behavioural profile as the DP/DTT-DMH targeted animals. Nevertheless, future experiments must definitively exclude a contribution of the LS to these behavioral effects.

Freezing behavior was scored automatically with Med Associates VideoFreeze software as previously described [136,137]. Due to interference of the patch cords with automated freezing counts, in all trials with patch cords freezing was hand-scored and cross-checked by an experimenter blind to experimental conditions.

### Perfusions and sectioning

Mice were euthanized 90 minutes after the fear memory retrieval test. Mice were injected with avertin (250mg/kg, i.p.) to induce deep anesthesia. Once unresponsive to tail and toe pinches, mice were transcardially perfused with 0.1M phosphate-buffered saline (PBS), followed by cold 4% paraformaldehyde fixative (PFA). The brains were extracted and stored in PFA overnight at 4°C. Brains were sectioned at 50µm (1 in 6 series) in the coronal plane with a vibratome (model VT1000, Leica). Sections were stored at -20°C in a cryoprotectant solution comprised of 60% glycerol and 0.01% sodium azide in 0.1M PBS.

### Serum corticosterone

A separate cohort of mice with no previous behavioral testing history (n=5 per group) was used to determine the effect of DP/DTT manipulations on serum corticosterone. The DP/DTT was injected with mCherry, hM4D, or hM3D viral constructs using identical parameters to those described earlier. After a 2-week recovery period, mice were weighed and acclimated to a procedure room for at least one hour. Mice were injected with C21 (2mg/kg, 0.5mg/mL, i.p.) and immediately returned to their home cage. Mice remained in their home cage for 90 minutes undisturbed and were then euthanized as described above. Immediately prior to perfusion, blood was extracted from the heart (∼0.3 to 0.5mL) with a 23-gauge needle attached to a 1mL syringe. The blood was immediately transferred to a 1.5mL Eppendorf tube and allowed to coagulate at room temperature for 30 minutes. Blood samples then underwent centrifugation to isolate serum (10,000 rpm, 10 minutes). The isolated serum was extracted with a 1mL pipette, transferred into a fresh microcentrifuge tube, and flash frozen with liquid nitrogen. Samples were stored at -20°C until use. Serum corticosterone was evaluated with a colorimetric competitive enzyme immunoassay kit (#ADI-900-097, Enzo Life Sciences) following the manufacturer’s instructions. Briefly, serum samples (1 in 40 dilution) were run in duplicate and read at 405nm with 570nm correction on a Biotek Synergy Neo2 Multi-Mode Plate Reader (Agilent) and analyzed using Biotek Gen5 software (version 3.08, Agilent).

### Immunofluorescence

Immunofluorescence staining was performed on free-floating sections. Sections were incubated with any of the following primary antibodies: polyclonal rabbit anti-mCherry (1:2000, catalog #ab167453, Abcam; RRID: AB_2571870), polyclonal chicken anti-GFP (1:3000, catalog #13970, Abcam; RRID: AB_371416), polyclonal rabbit anti-GluR2/3 (1:75, catalog # AB1506, Millipore; RRID: AB_90710), monoclonal mouse anti-GAD67 (1:250, catalog # MAB5406, Millipore; RRID: AB_2278725) overnight at 4°C on a rotary shaker under gentle agitation. On the following day, sections were washed in 0.1M PBS and then incubated with the following secondary antibodies: goat anti-rabbit Alexa Fluor 568 (1:500, catalog #A11011, Thermo Fisher Scientific; RRID: AB_143157), goat anti-mouse Alexa Fluor 633 (1:200, catalog # SAB4600138, Millipore), goat anti-chicken Alexa Fluor 488 (1:500, catalog # A11039, Thermo Fisher Scientific; RRID: AB_2534096), for 2hrs at room temperature. Hoechst 33342 (1:1000 diluted in 0.1M PBS, Thermo Fisher Scientific) was used as a counterstain. Sections were then rinsed in 0.1M, mounted onto gelatin-coated slides, and allowed to air dry for 30 minutes. The mounted sections were then coverslipped with Citifluor anti-fade mounting medium (catalog #17970, Electron Microscopy Sciences) or permafluor antifade medium (catalog # TA030FM, Fisher Scientific) and stored in a microscope slide storage book at 4°C until imaging.

To determine whether C21 modified DP/DTT activity across treatment groups, we processed a subset of PFC sections for the immediately early gene c-Fos. We chose sections from mice used in the serum corticosterone experiment because they did not undergo any behavioral testing which could influence patterns of c-Fos expression. Immunofluorescence staining for c-Fos was performed as previously described [136]. Sections were rinsed in 0.1M PBS and then incubated in 1% H_2_0_2_ in 0.1M PBS to quench endogenous peroxidase activity. Sections were incubated overnight at 4°C with monoclonal rabbit anti c-Fos (1:8000, catalog #226 008, Synaptic Systems; RRID: AB_2891278) and monoclonal rat anti-mCherry (1:3000, catalog #M11217, Thermo Fisher Scientific; RRID: AB_2536611) diluted in 0.1M Tris-buffered saline and 0.5% (w/v) Roche Blocking Reagent (catalog #11096176001, Sigma-Aldrich). On the following day, sections were rinsed in 0.1M PBS and incubated in goat anti-rat Alexa Fluor 568 secondary antibody (1:500, catalog #A11077, Thermo Fisher Scientific; RRID: AB_2534121) for 1 hr. Sections were then rinsed in 0.1M PBS and incubated in donkey anti-rabbit horseradish peroxidase- conjugated secondary antibody (1:500, catalog #711-036-152, Jackson ImmunoResearch; RRID: AB_2340590) for 1hr. Tyramide signal amplification was then performed with a fluorescein tyramide conjugate (1:400) diluted in 0.1M borate buffer containing 0.01% H_2_0_2_. Hoechst 33342 was used as a counterstain using the same parameters described above. Sections were then mounted onto gelatin-coated slides, air dried, and coverslipped with Citifluor anti-fade mounting medium as described above.

### Image acquisition & quantification

Images were acquired on a Nikon Eclipse Ni-U epifluorescence microscope connected to a computer running NIS-Elements software (v 5.11.03, Nikon). Immunofluorescence signal was visualized with a LED illumination system (X-Cite 120 LED Boost, Excelitas Technologies) and 10x Plan-Apochromat objective. Images were captured with a Nikon DS-Qi2 digital camera. A total of 3 images per mouse were used to evaluate viral expression in the DP/DTT for the mCherry, hM4D-mCherry, and hM3D-mCherry cohorts. We evaluated viral expression via the mCherry tag in sections located approximately +1.9mm to +1.7mm anterior-posterior (relative to bregma). Photomicrographs were superimposed on plates from the Paxinos Mouse Brain atlas and the spread of viral expression was drawn using the polygonal tool in Adobe Photoshop (version 24.1). Viral expression area for each mouse was represented using a semi-transparent color setting, as this helped visualize the areas with the most concentrated viral expression and viral spread across subjects.

For the DP/DTT TRAP neurochemical analysis, images were acquired using a Leica Stellaris 5 laser scanning confocal microscope. Confocal 1um Z-stack images of the DP/DTT were obtained and manually counted using LASX Office (version 1.4.5 27713) software (Leica).

Quantification of c-Fos expression in the DP/DTT was performed with ImageJ Software (version 1.53e) using procedures described previously [136]. Briefly, a minimum of 3 PFC sections per subject were used to count the number of c-Fos immunoreactive cells in the DP/DTT. The average number of cells per section was calculated by summing the total number of c-Fos+ cells across both hemispheres and dividing by the number of sections that were analyzed for each subject.

All figures were made in Adobe Photoshop. When brightness and/or contrast adjustments were made, these changes were applied to all photomicrographs.

### Slice electrophysiology

We used slice electrophysiology to confirm the effect of the DREADD agonist C21 on hM3D- and hM4D- transfected neurons in the DP/DTT and the effect of optical stimulation on ChR2 and GFP transfected DP/DTT axon terminals projecting to neurons in the DMH. Acute brain slices containing the mPFC were prepared from adult male and female C57BL/6J mice at least 14 days after AAV injection. For optogenetic experiments, electrophysiology recordings were conducted at least 1 week after the animals underwent behavioral testing.

Mice were deeply anesthetized with isoflurane and decapitated. The brain was quickly removed and placed in ice-cold sucrose based cutting solution containing the following (in mM): 180 sucrose, 2.5 KCl, 1.25 NaH_2_PO_4_, 25 NaHCO_3_, 1 CaCl_2_, 2 MgCl_2_, 2 Na^+^ pyruvate and 0.4 L-Ascorbic acid with 95% O_2_ and 5% CO_2_. The brain was cut in the coronal plane (300 µm thick) using a vibratome (VT1000S, Leica). Brain slices containing the mPFC were allowed to recover for 30 min in a holding chamber containing a recovery solution at 30 °C, made up of 50% sucrose-based cutting solution and 50% artificial cerebrospinal fluid (ACSF). The ACSF was composed of (in mM): 120 NaCl, 2.5 KCl, 1.25 NaH_2_PO_4_, 25 NaHCO_3_, 11 glucose, 2 CaCl_2_, 1 MgCl_2_, 2 Na^+^ pyruvate and 0.4 L-Ascorbic acid with 95% O_2_ and 5% CO_2_. Slices then underwent an additional 30 min recovery in regular ACSF at room temperature before patch-clamp recordings. After recovery, individual slices were transferred to a submersion chamber, perfused at 2 mL/min with ACSF at room temperature, and mounted on an upright microscope (Eclipse FN1, Nikon) that was equipped with a water-immersion long-working distance objective (40x), epifluorescence filters, and an infrared video camera [DAGE IR-2000].

For DREADD experiments, current-clamp recordings were obtained from mCherry+ pyramidal neurons in layers 2/3 or 5 of the DP/DTT with borosilicate pipettes (3–5 MΩ) that were filled with intracellular solution containing the following (in mM): 126 K D-Gluconate, 5 KCl, 10 HEPES, 4 MgATP, 0.3 NaGTP, 10 Na- phosphocreatine. Baseline data was acquired by measuring resting membrane potential (RMP) and the number of action potentials evoked by a long (1s) depolarizing current pulse. Somatic current pulses were calibrated in the baseline period to trigger approximately 3-4 action potentials (hM3D-mCherry^+^) or 7-8 action potentials (hM4D-mCherry^+^ pyramidal neurons). After 5 minutes of stable neuronal discharge induced by a somatic depolarization, C21 (1 µM, HelloBio) [150] was bath applied for at least 10 minutes, with the last 5 min being compared to the baseline period.

For optogenetic experiments, voltage clamp recordings were obtained from DMH neurons with borosilicate pipettes (3-5 MΩ) filled with intracellular solution containing the following (in mM): 130 Cs- methanosulfonate, 10 HEPES, 0.5 EGTA, 8 NaCl, 4 Mg-ATP, 0.4 Na-GTP, 10 Na-phosphocreatine, 1 N- ethyl lidocaine (QX-314). DP/DTT axon terminals were stimulated using a TTL-pulsed microscope objective-coupled light-emitting diodes (460 nm, ∼1.36 mW/mm2; Prizmatix). Evoked optical EPSCs and IPSCs were recorded at -70mV and 0mV from both ChR2 and GFP control transfected animals, respectively. All light-induced stimulation was conducted at 0.1 Hz to avoid inducing synaptic plasticity.

Electrophysiological data was acquired using a MultiClamp 700B amplifier (Molecular Devices). Signals were low-pass filtered at 2kHz and digitized at 20 kHz using Digidata 1550A and pClamp10 software (Molecular devices). Series resistance of the recorded neurons was regularly monitored and only cells with stable series resistance (less than 20% change of the initial value) and stable holding current were included in the experimental data.

### Statistical analysis

All results are presented as the mean ± standard error of the mean (SEM). Statistical comparisons were made in Prism 9.1 (GraphPad) with statistical significance (*p*<0.05) denoted on all graphs with an asterisk. Non-statistically significant results are reported in the figure legends due to space limits. All data were analyzed for sex differences, with non-statistically significant results reported below. To facilitate interpretation of the data, and in the absence of major differences within and across tasks, we pooled male and female mice, with individual data points presented as squares (males) and circles (females) for transparency.

For within subject comparisons, a paired t-test was used. Comparisons of independent groups were made using a one-way ANOVA, followed by Tukey’s Post Hoc test when appropriate. Parametric data with multiple comparisons were analyzed using a two-way repeated measures ANOVA, followed by Tukey’s or Šídák’s Post Hoc test with corrections for multiple comparisons. Normality of parametric datasets were confirmed by the D’Agostino and Pearson normality test (Prism 9.1).

### Sex differences

Potential sex differences were examined for each dataset using two-way ANOVAs followed by Šídák’s multiple comparisons test, and reported in the main text when significant. To facilitate interpretation of the data, and in the absence of major differences within and across tasks, we pooled male and female mice for each dataset, with individual data points presented as squares (males) and circles (females). For the OFT, we found no effect of sex on the time spent in the center zone (Two-way ANOVA, F_1,35_=1.065, *p*=0.309), outer zone (F_1,35_=0.553, *p*=0.462), or total distance travelled (F_1,35_=2.351, *p*=0.134). For the EPM, we found no sex differences in the time spent in the open arms (*F*_1,35_=3.069, *p*=0.009), time spent in the closed arms (*F*_1,35_=1.928, *p*=0.174), open arm entries (*F*_1,35_=0.386, *p*=0.538), or closed arm entries (*F*_1,35_=0.988, *p*=0.327). We found no sex differences in the TST (Two-way ANOVAs, time immobile *F*_1,35_=1.712, *p*=0.199; longest immobility bout *F*_1,35_=0.061, *p*=0.807; number of mobility-immobility transitions *F*_1,35_=2.044, *p*=0.162). In the FST, hM3D females spent significantly less time immobile than hM3D males and hM4D females (Two-way ANOVA, F_1,35_=6.561, *p*=0.015; Sidak’s multiple comparisons test, hM3D females versus hM3D males *p*=0.0093; hM3D females versus hM4D females *p*=0.0482; mCherry male versus female, *p*=0.264; hM4D male versus female, *p*=0.998). We did not observe any sex differences in the longest bout of immobility (F_1,35_=2.696, *p*=0.107), or the number of mobility-immobility transitions (F_1,35_=3.369, *p*=0.075) in the FST. In auditory fear acquisition, while we found a significant main effect of sex on average freezing to tone (Two-way ANOVA, F_1,35_=7.786, *p*=0.009). Post-hoc tests found that female hM4D mice froze significantly more than male hM3D mice (p=0.0479), but no other sex differences were observed (all *p* values >0.110). There were no sex differences in average tone freezing (Sidak’s multiple comparisons test, average tone freezing: mCherry *p*=0.126, hM3D *p*=0.351, hM4D *p*=0.505). We also failed to see sex differences in baseline freezing (two-way ANOVA, F_1,35_=0.179, *p*=0.675). We found no sex differences in auditory fear memory retrieval (Two-way ANOVA, average baseline freezing F_1,35_=0.397, *p*=0.533, average tone freezing F_1,35_=0.075, *p*=0.786).

Overall, we did not see sex differences in baseline OFT or EPM behavior. While studies in rats show consistent sex differences [151] in the EPM [151–157] and OFT [98,100,101,103,104 but see 105,106] which are often replicated in mice [160–164], several mouse studies show reversals or no sex differences [151,165–168], particularly in the OFT [151,160,168,169]. Although our lack of sex differences in the OFT replicates similar findings in C57BL6 mice [160,168,169], we unexpectedly found sex differences only in hM4D mice in the EPM. Similarly, our sex differences in the TST were only found among DREADD injected animals. While we cannot explain the lack of sex differences in the matched controls in these experiments, factors such as age [170,171] and estrous cycle [172,173] are known to modulate sex differences in affective behavior, and could have contributed to our findings. Although we have not seen major sex differences in the behaviors explored here, one study reported sex-specific increased recruitment of the DP in female mice following auditory fear conditioning [174]. Due to the well documented sex and gender differences in the prevalence of anxiety and posttraumatic stress (PTSD) disorders [175–177], as well as in sympathetic and autonomic function [178–181], this raises the intriguing possibility of sex differences in the long-term consequences of stress-induced activation of DP pathways.

